# The transcription factor GABPA is a master regulator of naïve pluripotency

**DOI:** 10.1101/2024.11.11.623003

**Authors:** Chengjie Zhou, Meng Wang, Chunxia Zhang, Yi Zhang

**Author notes:** These authors contributed equally to this work (Chengjie Zhou, Meng Wang, Chunxia Zhang). State Key Laboratory of Molecular Developmental Biology, Institute of Genetics and Developmental Biology, Chinese Academy of Sciences, Beijing, China (Chunxia Zhang).

## Abstract

The establishment of naïve pluripotency is a continuous process starting with the generation of inner cell mass (ICM) which then differentiating into epiblast (EPI). Recent studies have revealed key transcription factors (TFs) for ICM formation, but which TFs initiate EPI specification remains unknown. Here, using a targeted rapid protein degradation system, we show that GABPA is not only a regulator of major ZGA, but also a master EPI specifier required for naïve pluripotency establishment by regulating 47% of EPI genes during E3.5 to E4.5 transition. Chromatin binding dynamics analysis suggests that GABPA controls EPI formation at least partly by binding to the ICM gene promoters occupied by the pluripotency regulators TFAP2C and SOX2 at E3.5 to establish naïve pluripotency at E4.5. Our study not only uncovers GABPA as a master pluripotency regulator, but also supports the notion that mammalian pluripotency establishment requires a dynamic and stepwise multi-TFs regulatory network.

## Introduction

The development of mouse embryos starts with maternal-to-zygotic transition (MZT), which is accompanied by maternal RNA degradation and zygotic genome activation (ZGA) ^1, 2^. After a few rounds of cleavage, the totipotent embryos go through the first cell lineage differentiation to generate inner cell mass (ICM) and trophectoderm (TE) at E3.5 ^3, 4^. Subsequently, the ICM cells go through the second cell lineage differentiation to generate epiblast (EPI) and primitive endoderm (PrE) at E4.5 ^5^.

Transcription factors (TFs) play important roles in cell lineage specification and pluripotency acquisition. At E3.5, the ICM cells express both pluripotency factors such as *Oct4*, *Sox2* and *Nanog*, and PrE factors such as *Gata4* and *Gata6* ^6^. After completion of the second cell fate specification at E4.5, the expression of lineage-specific TFs become restricted with *Gata4* and *Gata6* confined to the PrE, while *Nanog* and *Sox2* restricted to the EPI, marking the establishment of naïve pluripotency in EPI ^6^. Given that embryonic stem cells (ESCs) cannot be fully established from E3.5 ICM ^7^, and E3.5 ICM cells have different transcriptome and chromatin accessibility from that of the E4.5 EPI ^8^, E3.5 ICM is considered to be at a ‘prepluripotency’ state ^8^. Recent studies indicate that the transcription factors NR5A2 and TFAP2C mediate totipotency to pluripotency transition by activating prepluripotency genes ^9^. However, neither of them is responsible for activating naïve pluripotency genes ^10^. Moreover, although NANOG and SOX2 are essential for maintaining the naïve pluripotency state, they are not required for initiating the naïve pluripotency *in vivo* ^11, 12^. Thus, the TFs that drive ICM to naïve pluripotency transition remain elusive.

Identification of the TFs regulating pluripotency establishment *in vivo* is hindered by their potential roles in earlier developmental stages. Thus, conventional TF knockout mouse models could not separate their potential roles in regulating totipotency or ICM formation from that in regulating EPI formation. The development of targeted protein degradation system like AID ^13^ and dTAG ^14, 15^ enable rapid degradation of target proteins at a specific time-window, making the study of gene function at a specific developmental stage possible. In this study, by generating dTAG mice and combined with RNA-seq and low-input CUT&RUN assay, we identified and demonstrated that the transcription factor GA repeat binding protein alpha (GABPA), encoded by the minor ZGA gene *Gabpa*, plays an essential role in naïve pluripotency establishment *in vivo*. Our results revealed that GABPA is not only important for major ZGA, but also critical for ICM to naïve pluripotency transition by regulating a large set of pluripotency genes through binding to their promoters. Importantly, we found that during ICM to EPI transition, the decreased binding of TFAP2C and SOX2 at the promoters of certain ICM genes concomitant with the increased binding of GABPA at these gene promoters, indicating a switch in the key TFs that regulate pluripotency genes expression. These results support a dynamic and stepwise regulatory model for naïve pluripotency establishment during pre-implantation development.

## Results

### Identification of GABPA as a potential pluripotency regulator

To identify candidate TFs potentially involved in ICM to naïve EPI differentiation, we performed integrative analyses of public RNA-seq and ATAC-seq datasets from mouse pre-implantation embryos ^16–19^. Compared to OCT4, NANOG, and SOX2 binding motif, GABPA binding motif is highly enriched in the open promoters of E4.5 ICM (EPI + PrE), suggesting GABPA may have a role in regulating E4.5 ICM formation (Fig. 1a). Consistent with a previous study ^16^, GABPA binding motif is not enriched in distal open chromatin (Fig. 1a). In addition, single-cell gene expression correlation analysis between various TFs and the ICM genes or TE genes (Supplementary Table 1) revealed that *Gabpa* expression positively correlates with expression of ICM genes in blastocysts, particularly at the late blastocyst stage similar to that of *Oct4* and *Nanog* (Fig. 1b), indicating potential involvement of *Gabpa* in pluripotency regulation. Furthermore, *Gabpa* transcription starts from zygote to early 2-cell stage, and reaches the highest at late 2-cell and 4-cell, and then become differentially expressed in ICM and TE at blastocyst (Fig. 1c). Immunostaining indicated that GABPA can be detected from zygote to blastocyst (Fig. 1d). These data are consistent with a potential role of GABPA in regulating pluripotency.

**Fig. 1.**
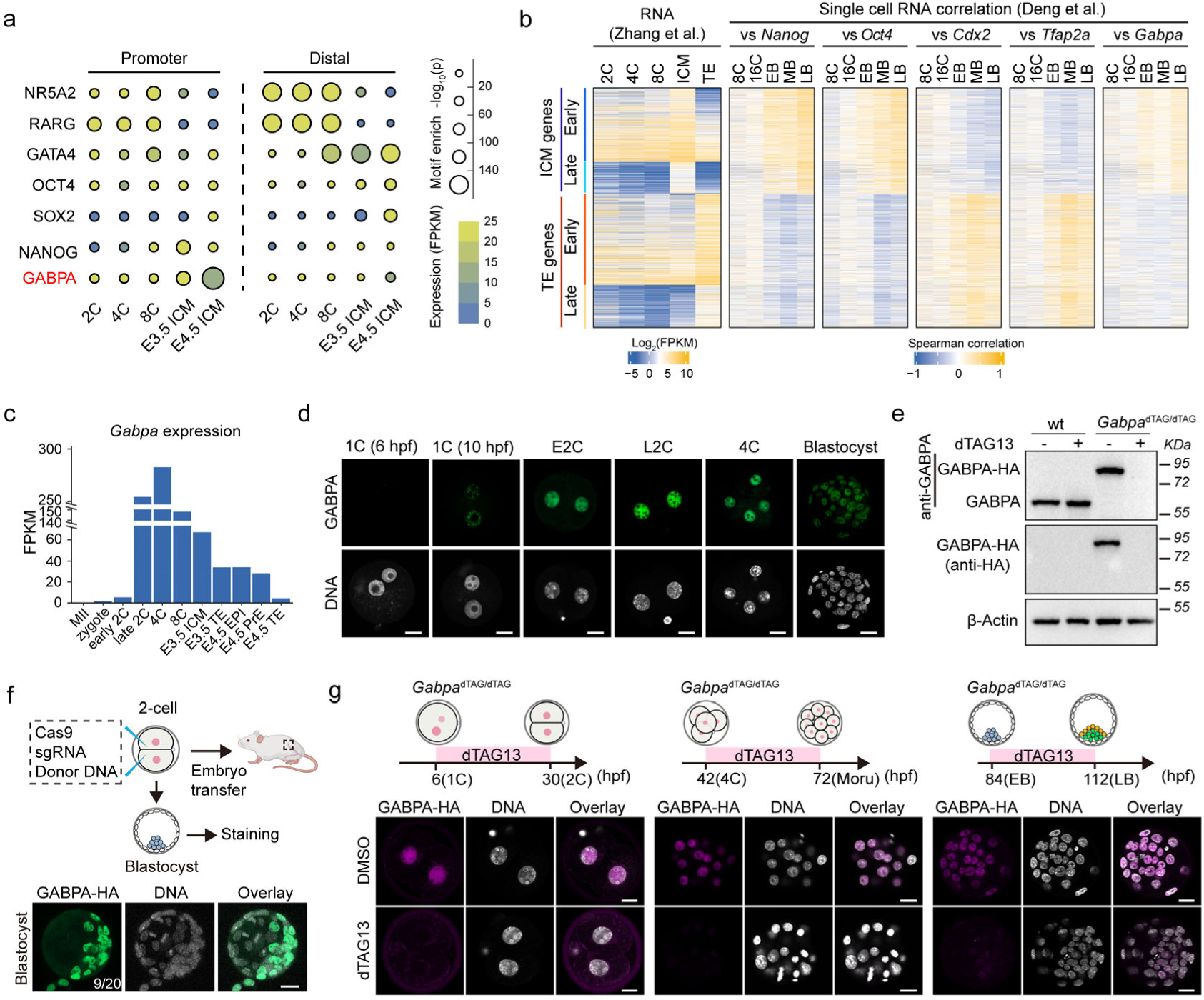
Identification of *Gabpa* as a potential pluripotency regulator and generation the *Gabpa*^dTAG/dTAG^ mice. (**a**) Enrichment of transcription factors motifs at promoter (left) and distal (right) ATAC-seq peaks (two biological replicates of each developmental stage were merged in this analysis) during mouse preimplantation development. (**b**) Expression correlation of ICM genes and TE genes with the expression of *Gabpa*, and other known ICM markers (*Nanog* and *Oct4*), TE markers (*Cdx2* and *Tfap2a*) at each stage of 8-cell (8C), 16-cell (16C), early blastocyst (EB), middle blastocyst (MB) and late blastocyst (LB) at single cell level. Early ICM/TE genes are genes with 8-cell expression (FPKM≥1), late ICM/TE genes are genes starting expression at E3.5 with no 8-cell expression (FPKM<1). Two biological replicates were merged. (**c**) *Gabpa* expression levels during mouse preimplantation development. (**d**) Immunostaining of GABPA (green) during mouse preimplantation development. DNA, Hoechst 33342. Scale bar, 20 μm. (**e**) Western blot confirming GABPA degradation with dTAG13 in ESCs. The blots were incubated with anti-GABPA and anti-HA, respectively. β-Actin was used as a loading control. (**f**) Strategy used for *Gabpa*^dTAG/dTAG^ mouse generation. HA staining indicates the dTAG knock-in cells in blastocyst. DNA, Hoechst 33342. Scale bar, 20 μm. (**g**) Immunostaining confirms GABPA degradation with dTAG13 treatment at different preimplantation developmental stages. anti-HA antibody was used for endogenous GABPA-HA fusion protein staining. DNA, Hoechst 33342. Scale bar, 20 μm. Experiments in **d**, **e**, **f** and **g** were repeated three independent times based on independent biological samples.

GABPA is a member of the ETS TFs family, which forms a tetrameric complex with GABPB to regulate the transcriptional activation ^20^. Previous studies have shown that *Gabpa* knockout resulted in embryonic lethal before blastocyst ^21^, but its role in cell fate specification is unknown. To assess its role in ICM and EPI specification, we utilized the dTAG system ^14^. Given the potential instability of dTAG fusion protein ^22^, we first tested GABPA dTAG system in mouse ES cells, which showed that the GABPA-FKBP-HA and the endogenous GABPA proteins are at similar levels (Fig. 1e and Extended Data Fig. 1a), indicating that the FKBP-HA fusion does not affect GABPA stability. Importantly, dTAG13 treatment resulted in complete GABPA-FKBP-HA degradation within 30 min (Fig. 1e). We thus proceeded to generate the mouse harboring the *Gabpa*^dTAG^ allele using the CRISPR technology (Fig. 1f and Extended Data Fig. 1a, b) ^23^. Importantly, addition of dTAG13 to the cultured embryos can efficiently degrade GABPA at zygote, 2-cell, 4-cell, morula, and blastocyst stages in both short- and long-time window treatments (Fig. 1g and Extended Data Fig. 1c, d). Moreover, dTAG13 treatment did not affect development of wide-type embryos (Extended Data Fig. 1e). Collectively, these results demonstrate the successful generation of a GABPA-dTAG mouse model.

### GABPA activates a group of major ZGA genes by binding to their promoters

To understand when and how GABPA degradation affects embryonic development, we performed dTAG13 treatment at different time windows. Given that *Gabpa* starts to express after fertilization (Fig. 1c), we confirmed that GABPA is a minor ZGA gene with a detectable protein level at 10 hrs post fertilization (hpf) (Fig. 1d). We therefore performed GABPA degradation starting at 6 hpf with continuous dTAG13 presence (Fig. 2a, #1), which caused most embryos arrest at 4-cell or 8-cell stage (Fig. 2b, #1, arrows), suggesting GABPA plays important roles before 4-cell. Since GABPA is highly expressed at late 2-cell stage (Fig. 1c, d), we asked whether GABPA regulates major ZGA. To this end, we treated the embryos with dTAG13 from 6 to 42 hpf and then washed out dTAG13 (Fig. 2a, b, #2). This treatment affected embryo development similarly to that when dTAG13 is continuously present from 6-112 hpf. Moreover, GABPA degradation after major ZGA (42-112 hpf) resulted in a much weaker phenotype (Fig. 2a, b, #3), suggesting GABPA plays an important role in ZGA.

**Fig. 2.**
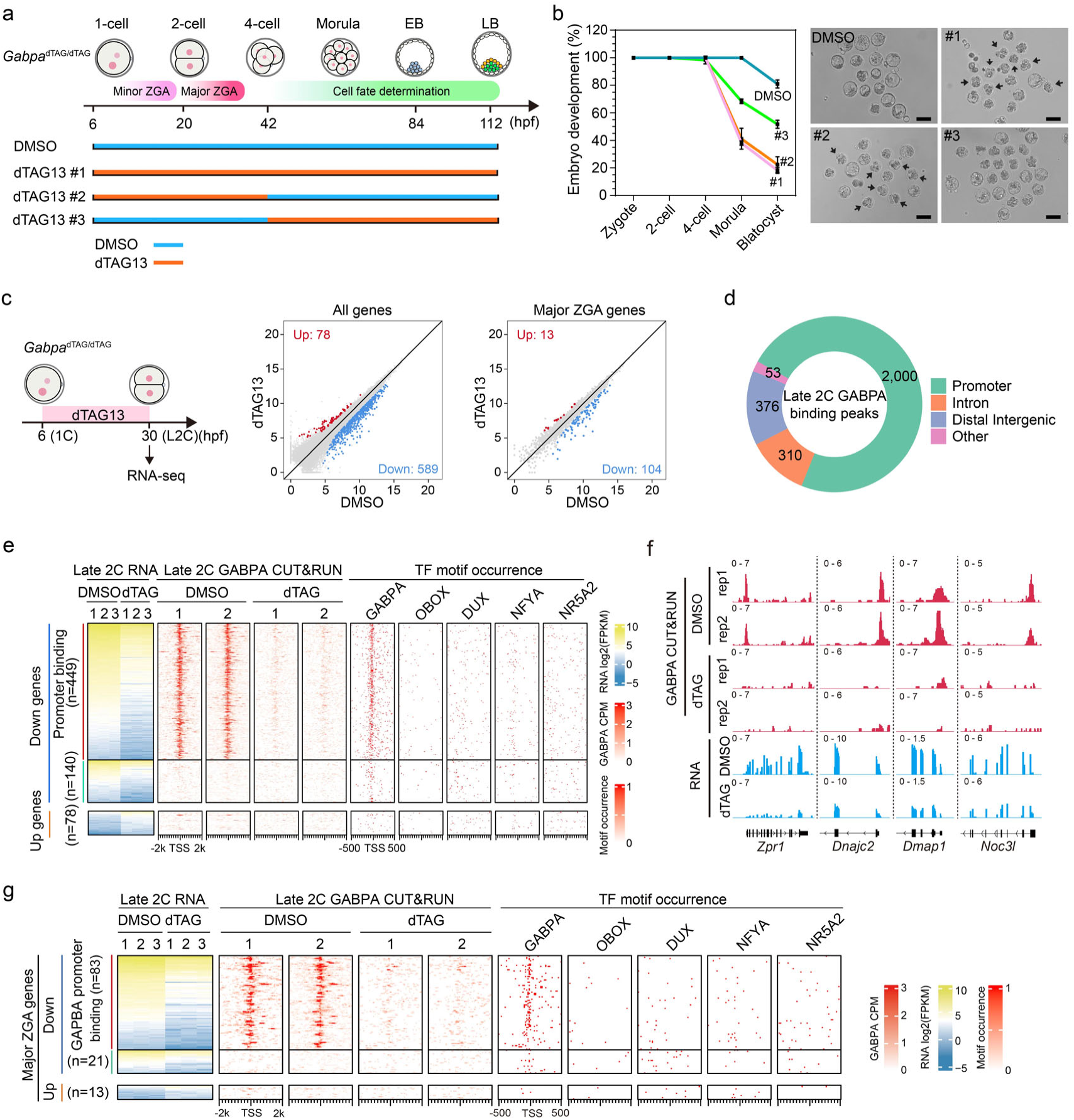
GABPA activates a group of major ZGA genes by binding to their promoters. (**a**) Diagram showing dTAG13 triggered GABPA degradation at different time window of mouse pre-implantation development. dTAG13 #1: dTAG13 treatment from zygote (6 hpf) to late blastocyst (112 hpf) stage; dTAG13 #2: dTAG13 treatment from zygote to 4-cell (42 hpf) stage (covers ZGA stage). dTAG13 #3: dTAG13 treatment from 4-cell to late blastocyst (after ZGA). (**b**) Embryo development rate (left) and representative images of blastocyst stage embryos (right) after GABPA degradation (DMSO, n=43; #1, n=50; #2, n=57; #3, n=54; n represents the total embryos from three independent experiments). Data are presented as mean values +/- SD. Black arrows indicate the 4-cell or 8-cell arrested embryos. Scale bar, 100 μm. (**c**) RNA-seq comparison of late 2-cell embryos treated with dTAG13 or DMSO from zygote (6 hpf) to late 2-cell (30 hpf). The x and y axis of the dot plots are log2 normalized counts from RNA-seq. (**d**) Genomic distribution of GABPA binding peaks generated by GABPA CUT&RUN at late 2-cell stage. (**e**) Heatmaps showing the differentially expressed genes (DEGs) at late 2-cell upon GABPA degradation, and the GABPA binding, TFs motif occurrence around the TSS of corresponding genes. The down-regulated genes were separated into two groups based on whether they have direct GABPA binding. (**f**) Genome browser examples of GABPA target genes at late 2-cell stage. (**g**) Heatmaps showing the differentially expressed major ZGA genes and GABPA binding profile in late 2-cell embryos after GABPA degradation. TFs motif occurrences around the TSS of corresponding ZGA genes are shown. The down-regulated genes were separated into two groups based on whether they have direct GABPA binding. Three biological replicates were used for RNA-seq analysis in panels **c**, **e** and **g**. The RNA-seq examples in panel **f** were showed as the pooled results of three biological replicates in each condition. Two biological replicates were used for CUT&RUN analysis in **d**, **e**, **f** and **g**.

To study GABPA’s role in regulating major ZGA, we treated the *Gabpa*^dTAG/dTAG^ embryos with dTAG13 from mid-zygote (6 hpf) to late 2-cell stage (30 hpf) and collected embryos at 30 hpf for RNA-seq (Fig. 2c and Extended Data Fig. 2a), which revealed 589 down-regulated and 78 up-regulated genes (Fig. 2c). GO analysis revealed that the down-regulated genes are involved in processes such as ribosome biogenesis, rRNA processing, etc. (Extended Data Fig. 2b, c), explaining the embryo arrest phenotype. In contrast, the up-regulated genes did not show GO term enrichment. Further analysis of the down-regulated genes identified 104 major ZGA genes, indicating GABPA has a role in activating these major ZGA genes (Fig. 2c and Supplementary Table 1, 2).

To determine whether GABPA directly binds to and regulates the down-regulated genes, we performed low-input CUT&RUN on GABPA ^24, 25^. We first tested GABPA CUT&RUN in mouse ESCs and found the use of 500 ESCs generated similar results as that of using 20k cells (Extended Data Fig. 3a, b). We then performed GABPA CUT&RUN using late 2-cell embryos (Extended Data Fig. 3b). Analysis of the CUT&RUN data indicated that most GABPA binding peaks are located in the nucleosome-depleted-regions (NDR) of promoters (Fig. 2d and Extended Data Fig. 3c). Interestingly, the GABPA binding regions were enriched for the GABPA motif but not the motifs of other murine ZGA regulators such as OBOX, DUX or NR5A2 ^26–28^, suggesting direct GABPA binding at these regions (Extended Data Fig. 3c). A clear GABPA binding signal around the transcription start sites (TSSs) of down-regulated genes (449 out of 589), but not up-regulated genes, was observed (Fig. 2 e, f). Importantly, most (83 out of 104) down-regulated major ZGA genes also showed direct GABPA promoter binding (Fig. 2f, g). The fact that GABPA degradation caused loss of promoter binding, resulted in down-regulation of most GABPA direct targets (Extended Data Fig. 3d, e), including some major ZGA genes (Fig. 2c, g), support that *Gabpa* plays an important role in activating these major ZGA genes by directly promoter binding.

### GABPA controls EPI specification by activating pluripotency genes

Next, we asked whether GABPA has a role in the first cell lineage specification. To avoid ZGA defect, we treated *Gabpa*^dTAG/dTAG^ embryos with dTAG13 starting from 4-cell, and collected early (84 hpf) and late blastocyst (112 hpf) for immunostaining (Fig. 3a). Neither ICM nor TE cell numbers in early blastocysts were affected by the treatment (Fig. 3b, c), suggesting GABPA is not required for the first cell lineage specification. However, for the late blastocyst, the EPI, but not the PrE, cells were dramatically decreased by the treatment (Fig. 3d, e). RNA-seq analysis of morula embryos revealed a limited transcriptome change upon the treatment (Extended Data Fig. 4a-c and Supplementary Table 3), consistent with minor role of GABPA from 4-cell to morula embryos. To avoid potential contribution from defects between 4-cell to early blastocyst, we started the treatment in early blastocyst when the first cell lineage specification has already finished, and then analyzed the effect on EPI and PrE at late blastocysts (Fig. 3f). A similar effect of this treatment to that of the treatment started at 4-cell embryos was observed (Fig. 3g, h), indicating that GABPA has a direct role in EPI specification.

**Fig. 3.**
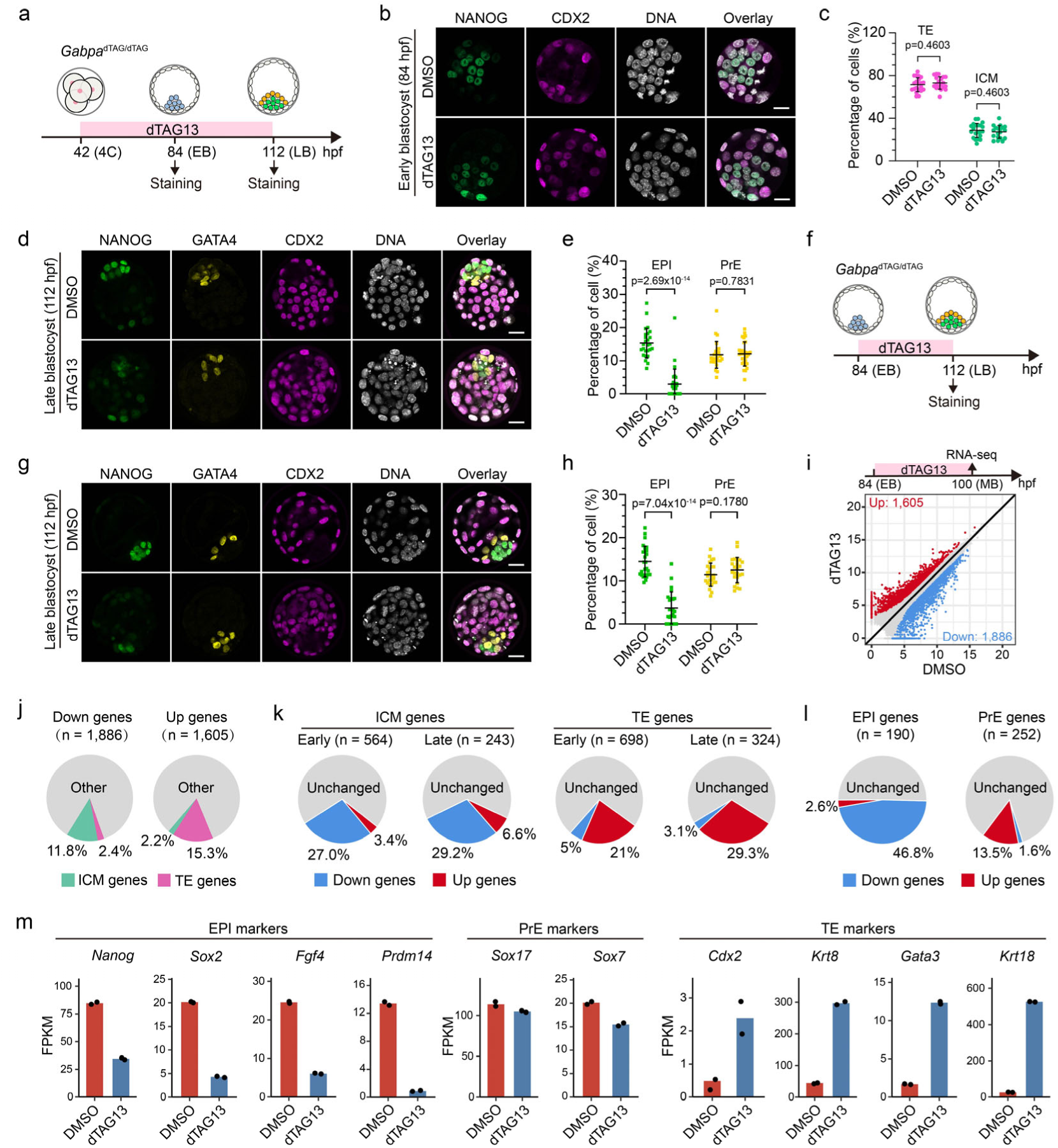
GABPA controls EPI specification by activating pluripotency genes. (**a**) Diagram showing the dTAG13 treatment after major ZGA and sample collection at early (84 hpf) and late (112 hpf) blastocyst. (**b**) Immunostaining of NANOG (green) and CDX2 (red) at early blastocyst with or without dTAG13 treatment. Scale bar, 20 μm. (**c**) Percentage of ICM and TE cells quantified based on panel b (DMSO, n=21; dTAG13, n=19. n represent total embryos of three independent experiments). ICM: NANOG^+^/CDX2^-^. TE: CDX2^+^. p values were calculated with Student’s t-test (two-sided). Data are presented as mean values +/- SD. (**d**) Immunostaining of NANOG (green), GATA4 (yellow) and CDX2 (red) at middle blastocyst stage with or without dTAG13 treatment. Scale bar, 20 μm. (**e**) Percentage of EPI and PrE cells quantifiedbased on panel d (DMSO, n=26; dTAG13, n=30. n represent total embryos of four independent experiments). EPI: NANOG^+^/GATA4^-^/CDX2^-^. PrE: GATA4^+^/ NANOG^-^/CDX2^-^. p values were calculated with Student’s t-test (two-sided). Data are presented as mean values +/- SD. (**f**) Diagram showing dTAG13 treatment from early to late blastocyst. (**g**) Same as panel d except the dTAG13 treatment time is as indicated in panel f. Scale bar, 20 μm. (**h**) Percentage of EPI and PrE cells quantified based on panel g (DMSO, n=26; dTAG13, n=24. n represent total embryos of three independent experiments). EPI: NANOG^+^/GATA4^-^/CDX2^-^. PrE: GATA4^+^/ NANOG^-^/CDX2^-^. p values were calculated with Student’s t-test (two-sided). Data are presented as mean values +/- SD. (**i**) DEGs from RNA-seq of E4.5 (late blastocyst) ICM with or without dTAG13 treatment. (**j**) Percentage of differential expressed ICM and TE genes. (**k**) Percentage of affected early/late ICM and TE genes after dTAG13 treatment. (**l**) Percentage of up- and down-regulated EPI and PrE genes after dTAG13 treatment. (**m**) Expression levels of EPI, PrE and TE marker genes with or without dTAG13 treatment.

To understand how GABPA regulates EPI specification, we performed RNA-seq. To capture the earlier molecular changes that lead to EPI defects, and to avoid potential confounding due to EPI and PrE cell ratio change, we collected ICM cells at the mid-blastocyst stage (100 hpf) when the EPI cell number has not yet shown difference between DMSO and dTAG13 treatment (Extended Data Fig. 4d-f). Transcriptome analyses revealed 1,605 up- and 1,886 down-regulated genes in response to GABPA degradation (Fig. 3i, Extended Data Fig. 4g, h and Supplementary Table 4). Some pluripotency factors have already expressed at E3.5 ICM when GABPA degradation starts (Extended Data Fig. 4d), despite EPI could form at middle blastocyst, these EPI cells are defective in their transcriptome due to GABPA degradation (Fig. 3i). Importantly, 223 of the 1,886 (11.8%) down-regulated genes belong to the ICM genes, while 246 of the 1,605 (15.3%) up-regulated genes belong to the TE genes (Fig. 3j). Further analysis revealed that 27% of the early ICM genes and 29.2% of the late ICM genes were down-regulated, while 21% of the early TE genes and 29.3% of the late TE genes were up-regulated by GABPA degradation, indicating that GABPA plays an essential role in the pluripotency establishment (Fig. 3k and Extended Data Fig. 4i).

We further found 46.8% of EPI genes were down-regulated (Fig. 3l, m, Extended Data Fig. 4j and Supplementary Table 1). In contrast, much smaller percentage of PrE genes (1.6%) were down-regulated, and importantly, most PrE marker genes, such as *Sox17* and *Sox7,* did not show significant change (Fig. 3l, m and Extended Data Fig. 4k), consistent with EPI but not PrE were affected upon GABPA degradation (Fig. 3d-h). Collectively, our data support GABPA determines EPI but not PrE specification by regulating EPI genes.

### GABPA regulates EPI gene expression by promoter and distal binding

To understand how GABPA regulates the ICM genes in E4.5 embryos, we performed GABPA CUT&RUN using E4.5 ICM cells (Extended Data Fig. 5a), which revealed that GABPA mainly occupies the promoter regions (Fig. 4a and Extended Data Fig. 5b). Motif analysis revealed that these regions are enriched for the GABPA motif, but not motifs of other lineage regulators including OCT4, SOX2 and NANOG, etc., indicating GABPA plays a direct role by binding to these regions (Extended Data Fig. 5b). Integrative analyses of the GABPA CUT&RUN and RNA-seq data revealed that almost half of the down-regulated genes (904 out of 1,886) were bound by GABPA, which is consistent with its motif enrichment in these promoters (Fig. 4b). In contrast, much fewer up-regulated genes are directly bound by GABPA (Fig. 4b). These results indicate that GABPA directly binds to promoters to activate these genes in E4.5 ICM.

**Fig. 4.**
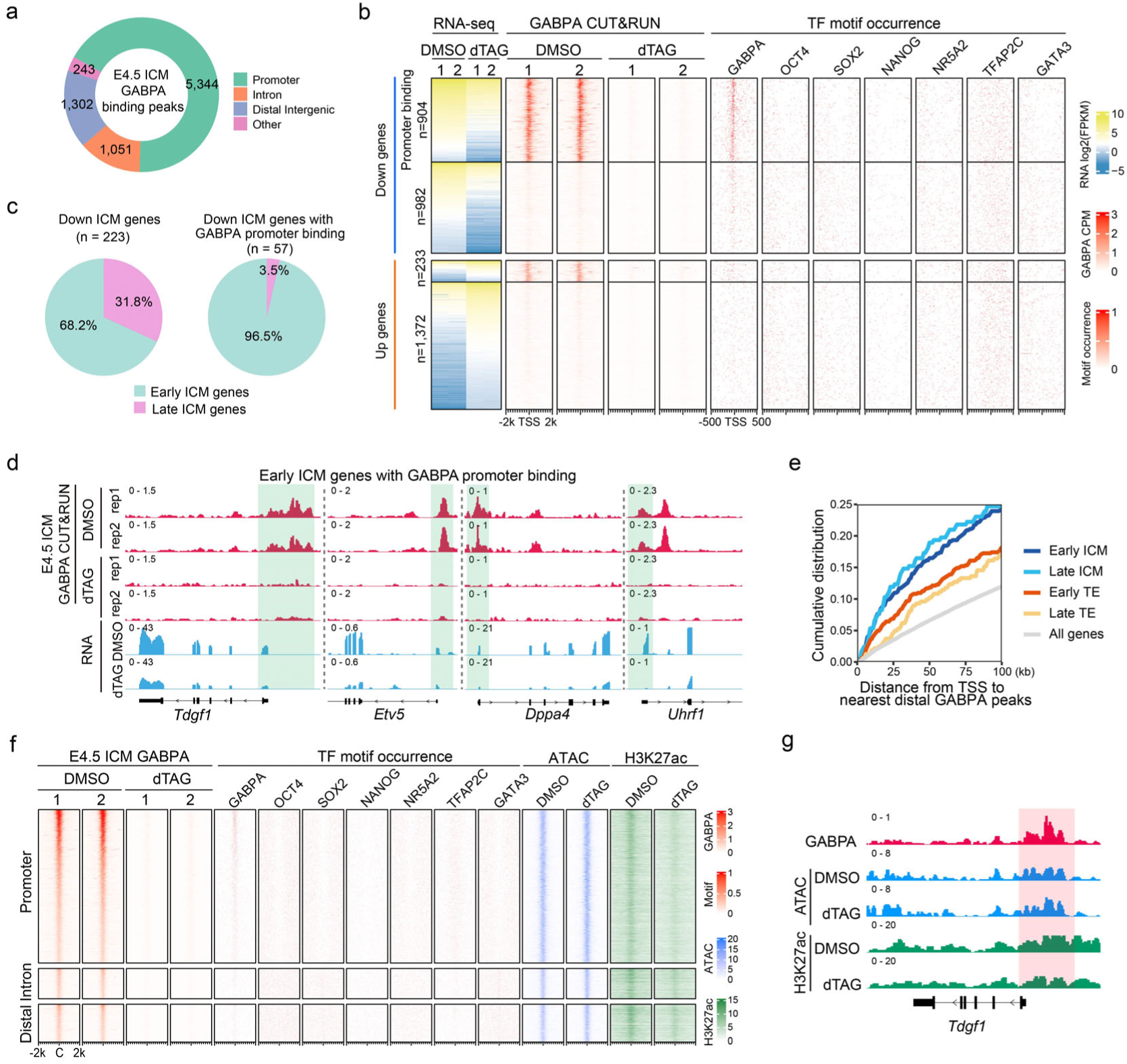
GABPA regulates EPI genes in E4.5 ICM by both promoter and distal binding. (**a**) Genomic distribution of GABPA binding peaks in E4.5 ICM based on CUT&RUN data. (**b**) Heatmaps showing the DEGs at E4.5 ICM after dTAG13 treatment, as well as the GABPA binding and several TFs motif occurrence around the TSS of the indicated gene groups. (**c**) Percentage of the down-regulated early and late ICM genes (left), and down-regulated ICM genes with GABPA promoter binding (right) at E4.5 ICM. (**d**) Examples of early ICM genes with GABPA promoter binding (shaded) at E4.5 ICM. (**e**) Cumulative distributions of the distance from TSS to the nearest distal GABPA peaks for each group of genes at E4.5 ICM. (**f**) Heatmap showing all the GABPA peaks at promoters, introns and distal regions in E4.5 ICM, and ATAC-seq and H3K27ac signals at these GABPA peak regions with or without dTAG13 treatment. C: peak center. (**g**) An example of genome browser view of GABPA binding (shaded), ATAC and H3K27ac signals in E4.5 ICM with or without dTAG13 treatment. All RNA-seq, GABPA CUT&RUN, ATAC-seq and H3K27ac CUT&RUN analysis were performed using two independent biological replicates. The pooled results for ATAC-seq and H3K27ac CUT&RUN are shown in panels **f** and **g**.

By separating E4.5 ICM genes into early and late ICM genes, we found that GABPA mainly bound to the promoters of early ICM genes (e.g., *Tdgf1*, *Etv5, Dppa4* and *Uhrf1*), but not late ICM genes (Fig. 4c, d). In addition to promoter, GABPA also exhibited distal binding, which were putative enhancer regions. We calculated the distance between TSS of ICM/TE genes and their nearest distal GABPA peaks and found that the TSS of the ICM genes were closer to the distal GABPA peaks than that of the TE genes (Fig. 4e), indicating a potential role of these distal GABPA bindings in regulating ICM genes. Indeed, we detected binding of GABPA to the known super-enhancer of ICM gene *Sox2* ^29^ (Extended Data Fig. 5c).

To investigate whether GABPA would affect chromatin accessibility and/or promoter/enhancer activity, we performed ATAC-seq as well as H3K27ac CUT&RUN in E4.5 ICM with or without GABPA degradation (Extended Data Fig. 5d). We found that GABPA degradation resulted in a widespread decrease of H3K27ac with little effect on chromatin accessibility (Fig. 4f, g and Extended Data Fig. 5e). This result indicates that while GABPA is not responsible for chromatin opening, it is important for the transcriptional activity of the bound genes. Previous reports of the interaction between GABPA and p300 ^30, 31^ in combination with the fact that GABPA degradation results in a decrease in H3K27ac level suggest that GABPA may activate its targets by recruiting the acetyltransferase p300.

### GABPA regulates a common sets of EPI genes in E4.5 ICM and naïve ESCs

Since E4.5 ICM is composed of EPI and PrE cells, it is technically challenging to obtain pure EPI cells to evaluate the role of GABPA in regulating EPI gene expression. To overcome this challenge, we used 2iESCs as they are believed to resemble the E4.5 EPI^32^. Since *Gabp*a knockout affects ESCs survival ^33^, we used the *Gabpa*^dTAG/dTAG^ ESCs treated with dTAG13 to evaluate the acute effect of GABPA degradation (Extended Data Fig. 6a). GABPA degradation was confirmed by Western blot analysis (Fig. 1e) and immunostaining (Extended Data Fig. 6b). RNA-seq revealed 2,265 down-regulated genes and 1,563 up-regulated genes in response to GABPA degradation for 24h (Extended Data Fig. 6c, d and Supplementary Table 5). Importantly, GABPA CUT&RUN analysis revealed that 1,238 (54.7%) of the down-regulated genes have GABPA promoter binding (Fig. 5a and Extended Data Fig. 6e). Similar to that in E4.5 ICM, GABPA binding motif was enriched in the GABPA peaks and its removal has little effect on ATAC-seq signals (Extended Data Fig. 6f, g). In contrast, the up-regulated genes have much fewer GABPA direct promoter binding (Fig. 5a). Comparative analysis confirmed that GABPA binding in E4.5 ICM and ESCs were highly similar (Fig. 5b), suggesting that GABPA may regulate the same set of genes in ESCs and E4.5 ICM. Indeed, analysis of the differentially expressed genes (DEGs) in E4.5 ICM and ESCs revealed significant overlap with 933 commonly down-regulated, and 356 commonly up-regulated genes in response to GABPA degradation (Fig. 5c). These results indicate that GABPA regulates a similar set of genes in E4.5 ICM and 2i ESCs.

**Fig. 5.**
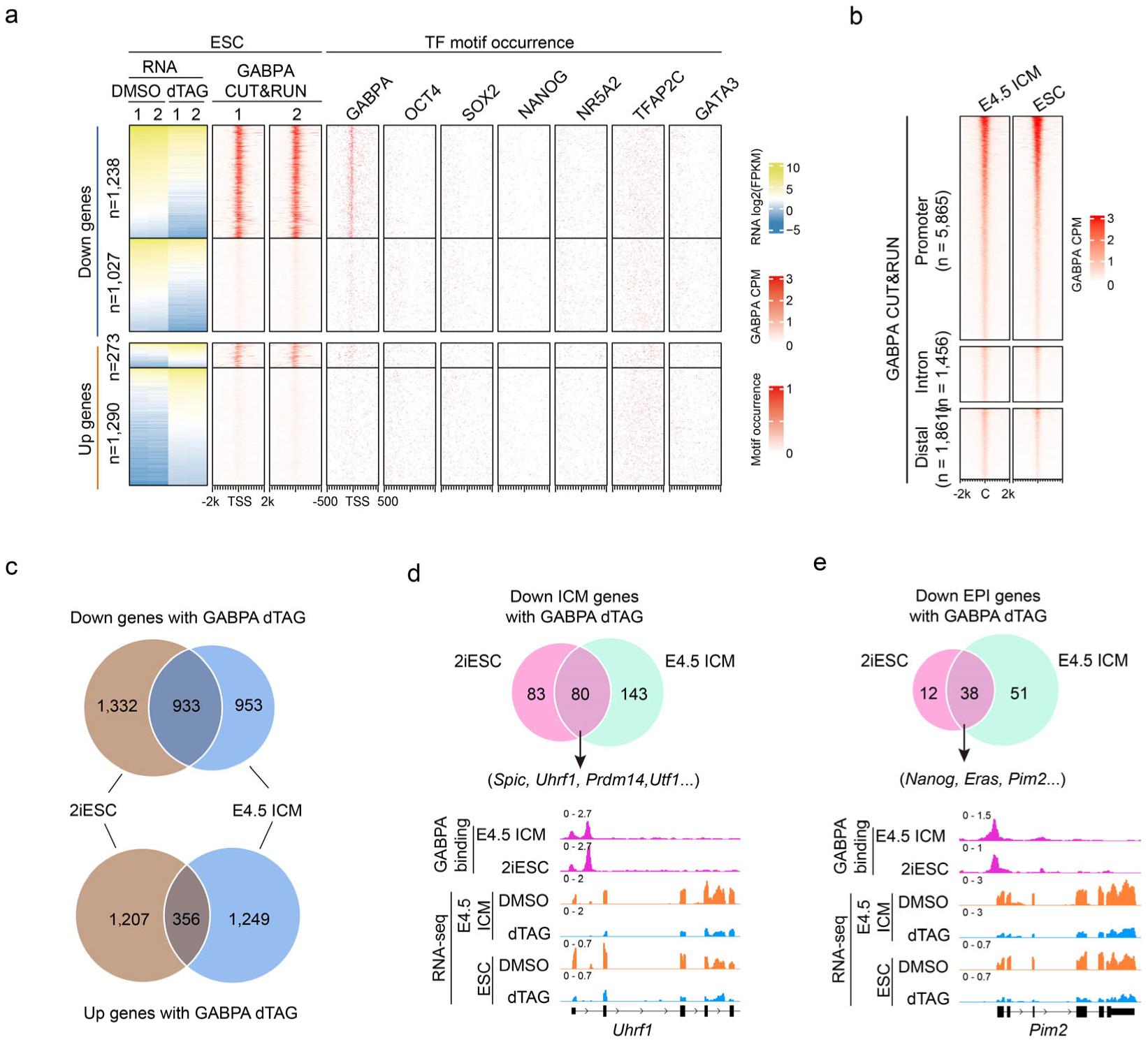
GABPA regulates a common set of EPI genes in E4.5 ICM and ESC. (**a**) Heatmaps showing the DEGs in 2iESCs after dTAG13 treatment for 24h, and the GABPA binding, GABPA motif occurrence around the TSS of the indicated gene groups. (**b**) Heatmaps showing similar GABPA binding peaks in E4.5 ICM and 2iESCs. Two replicates were merged for GABPA binding analysis in E4.5 ICM and ESCs. (**c**, **d**, **e**) Comparison of the DEGs (c), down-regulated ICM genes (d), and down-regulated EPI genes (e) between E4.5 ICM and 2iESCs after dTAG13 treatment. RNA-seq and GABPA CUT&RUN analysis were performed using two independent biological replicates. Panel **b**, **d** and **e** showed the pooled results of the two biological replicates in each condition.

We further found that 26.3% EPI genes were down-regulated and 12.7% PrE genes were up-regulated by GABPA degradation (Extended Data Fig. 6h), suggesting GABPA plays an important role in pluripotency regulation in ESCs. Further analysis of the down-regulated ICM and EPI genes revealed 80 ICM genes are in common (e.g., *Spic, Utf1, Prdm14* and *Utf1*) (Fig. 5d), including 60 early ICM genes and 20 late ICM genes (Extended Data Fig. 6i). The analysis also revealed 38 EPI genes in common (e.g., *Nanog, Eras* and *Pim2*) (Fig. 5e). Collectively, data from E4.5 ICM and 2iESCs support the notion that GABPA plays a crucial role in regulating naïve pluripotency both *in vivo* and *in vitro*.

### Stepwise pluripotency establishment controlled by TFAP2C/SOX2/GABPA

The first and second cell fate specification in mouse embryo occurs at E3.5 and E4.5 with pluripotency gene expression restricted within ICM and EPI, respectively ^34–36^. However, the expression of some ICM genes starts as early as the 2-cell stage (Fig. 1b). A recent study revealed the role of the transcription factors NR5A2 and TFAP2C in activating ICM genes at the 8-cell stage ^10^. However, the expression of *Nr5a2* and *Tfap2c* are respectively silenced at E3.5 ICM and E4.5 EPI (Extended Data Fig. 7a), indicating other transcription factors are responsible for the ICM gene activation at these stages. Our data indicate that GABPA plays such a role for EPI gene activation at E4.5.

To understand how GABPA participates in this process, we generated additional GABPA CUT&RUN dataset in 8-cell and E3.5 ICM (Extended Data Fig. 7b, c). Comparative analysis of GABPA binding profiles at 2-cell, 8-cell, E3.5 ICM and E4.5 ICM indicate strong promoter GABPA binding occurs at 2-cell stage and is maintained through E4.5 ICM (Fig. 6a and Extended Data Fig. 7d), which is consistent with the continuous expression of their target genes (Extended Data Fig. 7e). Weak promoter bindings at 2-cell became stronger at E4.5 ICM stage (Fig. 6a). Increased GABPA binding to promoters of ICM genes at E4.5 ICM likely contributes to GABPA’s stage-specific functions (Fig. 6a and Extended Data Fig. 7f).

**Fig. 6.**
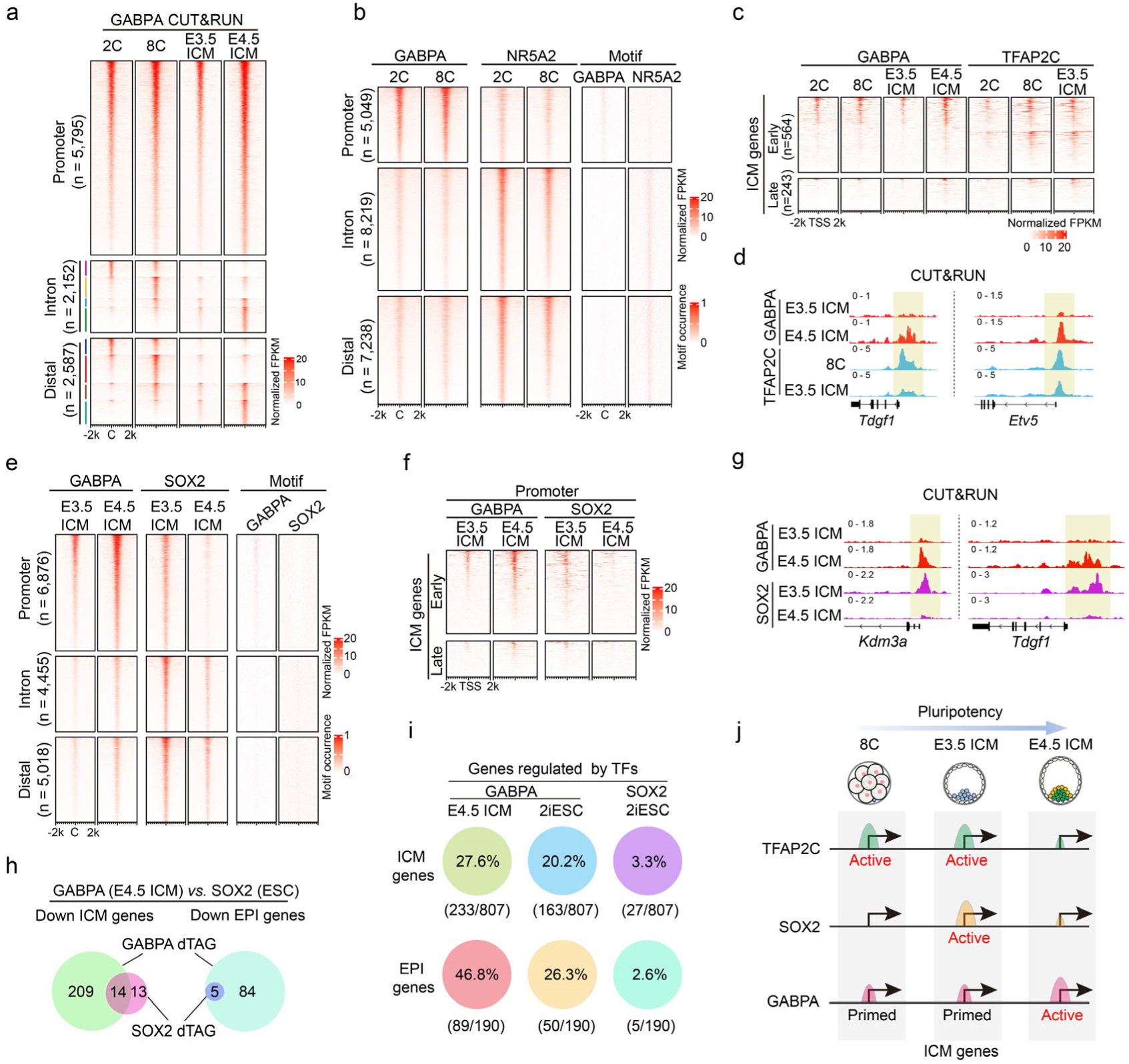
GABPA genomic binding dynamics during pluripotency establishment and its relationship with other pluripotency regulators. (**a**) Heatmaps showing the dynamics of GABPA genomic binding profile from 2-cell, 8-cell and E3.5 ICM to E4.5 ICM at promoter, intron and distal regions. C: peak center. (**b**) Comparison of GABPA and NR5A2 binding profiles and motif occurrence at promoter, intron and distal regions at 2-cell and 8-cell stages. (**c**) GABPA and TFAP2C binding profiles at promoters of early and late ICM genes in 2-cell, 8-cell and E3.5 ICM. (**d**) Genome browser view of two examples showing GABPA and TFAP2C binding dynamics. Promoter binding regions are shaded. (**e**) GABPA and SOX2 binding profiles at promoter, intron and distal regions in E3.5 ICM and E4.5 ICM. (**f**) GABPA and SOX2 binding profiles at promoters of early and late ICM genes in E3.5 ICM and E4.5 ICM. (**g**) Genome browser view of two examples showing the binding (shaded) of GABPA and SOX2 in E3.5 ICM and E4.5 ICM. (**h**) Comparisons of GABPA (E4.5 ICM) and SOX2 (ESC) affected ICM genes (left) and EPI genes (right) after dTAG13 treatments. (**i**) Percentage of ICM and EPI genes regulated by GABPA and SOX2 in E4.5 ICM and ESCs. (**j**) A model illustrating different TFs including TFAP2C, SOX2 and GABPA play major roles at different stages during the pluripotency establishment (from 8-cell stage to E4.5 ICM stage). CUT&RUN analysis in panels **a-g** were performed using two independent biological replicates, and pooled results were used.

Next, we analyzed the relationship between GABPA and NR5A2 by comparing their binding profiles at 2-cell and 8-cell. Interestingly, GABPA tends to bind to promoters, while NR5A2 preferentially binds to putative enhancers (Fig. 6b), suggesting that they have distinct mechanisms in gene regulation. We also compared the binding profiles of GABPA and TFAP2C at 2-cell, 8-cell and E3.5 ICM and found TFAP2C and GABPA co-occupy a portion of ICM genes promoters at 8-cell and E3.5 ICM (Extended Data Fig. 7g), which may explain why GABPA degradation at these stages does not show phenotype. Given that TFAP2C is not expressed in E4.5 EPI (Extended Data Fig. 7a) ^9^, while GABPA binding is increased from E3.5 ICM to E4.5 ICM, GABPA may take over TFAP2C’s function on their commonly occupied ICM gene promoters (Fig. 6c, d). To identify the commonly regulated genes taken over by GABPA, we compared the down-regulated genes in GABPA-dTAG E4.5 ICM with the down-regulated genes in *Tfap2c* maternal-zygotic KO (mz-KO) E3.5 ICM and identified 7 out of the 13 *Tfap2c* KO down-regulated ICM genes and all the 5 *Tfap2c* KO down-regulated EPI genes showed overlap (Extended Data Fig. 7h). Since *Tfap2c* mz-KO at 8-cell showed more down-regulated ICM and EPI genes than that in E3.5 ICM (Extended Data Fig. 7i), TFAP2C plays more important role in activating pluripotency genes at 8-cell than in E3.5, which is consistent with the observation that *Tfap2c* expression is nearly undetectable at E3.5 ICM (Extended Data Fig. 7a) ^9^.

*Sox2* starts to express at E3.5 ICM and plays an important role in regulating pluripotency (Extended Data Fig. 7a) ^8^. We thus compared the binding profiles of GABPA and SOX2 in E3.5 ICM and E4.5 ICM. We found a global increase in GABPA binding concomitant with a global decrease in SOX2 binding, especially on promoters from E3.5 ICM to E4.5 ICM (Fig. 6e). Consistently, GABPA binding on ICM gene promoters was enhanced, while SOX2 binding on ICM gene promoters was lost (Fig. 6f, g), indicating the regulation of ICM genes by GABPA was enhanced at E4.5 stage. To explore the possibility that GABPA takes over the role of SOX2 in activating their commonly regulated ICM genes at E4.5, we compared the down-regulated genes in GABPA-dTAG E4.5 ICM and *Sox2-*mzKO E3.5 ICM and found 18 out of the 55 *Sox2-*KO down-regulated ICM genes and 8 out of the 9 *Sox2-*KO down-regulated EPI genes showed overlap (Extended Data Fig. 7j), supporting the notion that GABPA takes over SOX2’s role on these ICM gene promoters in E4.5 ICM. To gain further support for this notion, we compared the down-regulated genes of GABPA-dTAG E4.5 ICM with that of the SOX2-dTAG 2iESCs. Although SOX2 degradation in 2iESC only down-regulated a small number of ICM (27) and EPI (5) genes, all the SOX2 down-regulated EPI genes were also down-regulated by GABPA and more than half of the SOX2 down-regulated ICM genes were also down-regulated by GABPA in embryo (Fig. 6h). 3 out of the 5 SOX2-dTAG down-regulated EPI genes and 7 out of the 27 down-regulated ICM genes were also down-regulated by GABPA degradation in ESCs (Extended Data Fig. 7k). Collectively, these data support our notion that TFAP2C and SOX2 are responsible for activating pluripotency genes at 8-cell and E3.5 ICM, respectively, while GABPA mainly activates pluripotency genes during E3.5 ICM to E4.5 EPI transition. GABPA takes over the role of TFAP2C and SOX2 at the promoter of pluripotency genes in E4.5 EPI in addition to activating its unique targets.

## Discussion

Zygotic genome activation and pluripotency acquisition are two of the most important events during preimplantation development. Understanding the molecular mechanisms underlying these events are not only important in the field of development, but also critical for regenerative medicine. Here we identified and demonstrated that a minor ZGA factor GABPA not only regulates major ZGA, but also plays a critical role in pluripotency establishment by activating a large group of pluripotency genes in E4.5 EPI. Despite the observation that GABPA KO down-regulates pluripotency factors *Nanog* and *Oct4* in ESCs ^33, 37^, its role as a key pluripotency regulator was not recognized due to the lack of direct genome binding data. Additionally, its KO lethality phenotype before blastocyst formation ^21^ prevented its role in pluripotency *in vivo* being addressed using the conventional KO mouse model. In this study, using the dTAG system, we reveal the developmental stage-specific function of GABPA during mouse pre-implantation development. Our data support GABPA plays a non-dispensable role in ZGA and serves as a master regulator of pluripotency establishment at E4.5 EPI.

Identification of TFs important for mammalian ZGA has been a hot topic in the past several years. Previous studies have been mainly focused on maternal factors that regulate mouse ZGA such as OBOX ^26^, NR5A2 ^28^, DUX ^27^, NFYA ^38^ and KLF17 ^39^, while the contributions of zygotic early transcribed genes are neglected although published work have showed that minor ZGA is required for major ZGA in mice ^40^. Here, we provide evidence demonstrating that the minor ZGA factor GABPA regulates a subset of major ZGA genes and pre-implantation development by directly binding to the promoter of 83 major ZGA genes to activate their transcription (Fig. 2g). Our study indicates that minor ZGA gene product can regulate major ZGA by directly binding to and activating major ZGA. Further studies of the other minor ZGA genes are warranted to fully understand the role of minor ZGA genes in regulating major ZGA and pre-implantation development.

Another important finding of this study is that we identify GABPA as a key TF driving the E4.5 EPI specification. We showed that GABPA degradation affect EPI, but not PrE formation (Fig. 3d-h), which could be explained by GABPA’s role in activating pluripotency genes including *Nanog* and *Sox2* (Fig. 3m). Different from *Nanog* and *Sox2*, which are exclusively expressed in EPI, *Gabpa* is expressed in both EPI and PrE. Such expression pattern raises an intriguing question about why GABPA selectively activates EPI genes, but not PrE genes (Fig. 3l, m). One possibility is that binding of GABPA to the pluripotency gene promoters requires another factor that is only expressed in EPI. Alternatively, PrE may express a protein that can mask GABPA binding to promoters. Future studies should confirm or refusal these possibilities.

Establishment of pluripotency is a continuous and complex process. Although the expression of pluripotency genes such as *Nanog*, *Sox2* and *Oct4* are critical for maintaining pluripotency ^41–44^, their role in pluripotency establishment has not been shown. At 8-cell and E3.5 ICM, TFs such as NR5A2, TFAP2C and SOX2 are believed to modulate the activity of these pluripotency genes ^8, 9^, they are unlikely have a major role in EPI specification as they are either not expressed in EPI or they only occupy the promoters of very few pluripotency genes in EPI. Our data indicate that GABPA can fulfil such a role as it regulates 46.8% and 26.3% of EPI genes in E4.5 and 2iESC, respectively (Fig. 6i). Although the lack of a dTAG mouse model for SOX2 prevented a direct comparison of the role of GABPA and SOX2 in EPI formation, a comparison of dTAG13-mediated degradation of GABPA or SOX2 in 2iESC showed a clear difference as GABPA degradation down-regulated many more ICM genes and EPI genes than that of SOX2 (Fig. 6i). Our study, together with previous studies, supports a stepwise model for naïve pluripotency establishment (Fig. 6j). At 8-cell, TFAP2C binds to gene promoters and enhancers to active pluripotency genes. At E3.5, with the decrease of TFAP2C expression in ICM, its role on pluripotency gene regulation decreases, while SOX2 starts to function. By E4.5, TFAP2C is no longer expressed in ICM (Extended Data Fig. 7a) ^9^, and the binding of SOX2 to the promoters of ICM genes also disappeared, while GABPA now occupies and activates almost half of the pluripotency gene in EPI. Our study thus not only identifies GABPA as an important TF for ZGA, but also demonstrates its crucial role as a master regulator of naïve pluripotency in E4.5 embryos.

## Acknowledgements

We thank Dr. Yota Hagihara and Dr. Qianying Yang for their valuable discussions on the project and comments on the manuscript. This project was supported by the NIH (R01HD092465) and the HHMI. YZ is an investigator of the Howard Hughes Medical Institute. This article is subject to HHMI’s Open Access to Publications policy. HHMI lab heads have previously granted a nonexclusive CC BY 4.0 license to the public and a sublicensable license to HHMI in their research articles. Pursuant to those licenses, the author-accepted manuscript of this article can be made freely available under a CC BY 4.0 license immediately upon publication.

## Author contributions

Y.Z. conceived and supervised the project. C-J.Z. and M.W. designed the experiments. C-J.Z. performed the experiments including embryo manipulation, RNA-seq, CUT&RUN, ATAC-seq, etc. M.W. performed all the bioinformatic analysis. C-X.Z. established the GABPA-dTAG mice and GABPA-dTAG ESCs and performed WB verification. C-J.Z., M.W. and Y.Z. wrote the manuscript. All authors interpreted the data and reviewed the manuscript.

## Methods

### Animals

All experiments were conducted in accordance with the National Institute of Health Guide for Care and Use of Laboratory Animals and approved by the Institutional Animal Care and Use Committee (IACUC) of Boston Children’s Hospital and Harvard Medical School (protocol number IS00000270-6). All mice were kept under specific pathogen-free conditions within an environment controlled for temperature (20-22°C) and humidity (40-70%), and were subjected to a 12-hrs light/dark cycle. Generation of *Gabpa*-dTAG knock-in mice was as described previously with some modification ^23^. Briefly, 2-cell embryo (20 hpf) were injected with *Gabpa* donor DNA (30 ng/μl), Cas9 mRNA (100 ng/μl) and sgRNA (50 ng/μl) using a Piezo impact-driven micromanipulator (Primer Tech, Ibaraki, Japan). Then 2-cell embryos were incubated in KSOM for 2 hrs before transferred into oviducts of pseudo-pregnant ICR strain mothers (Charles River). F0 chimera mice was backcross with wild-type C57BL/6J mice for at least two generation. Genotyping was performed using mouse tail lysed in lysis buffer (50 mM Tirs-HCl, 0.5% Triton and 400 μg/ml Proteinase K) at 55°C overnight. For F0 and F1 mice genotyping, the primers outside the homology arm are used. For genotyping of F2 and beyond, the inner primers are used. The primers are listed in Supplementary Table 6.

Droplet Digital PCR (ddPCR) was used for detecting the copy number of *Gabpa*-dTAG knock-in allele in F1 mice. Briefly, 250 ng purified DNA templates were digested by incubation with Haelll Enzyme (NEB) at 37°C for 1h, and then inactivated at 80°C for 5 mins. A final 30 ng DNA was used as the templates for PCR. *Fkbp* was used for knock-in detecting, *mRPP30* was used as a control. The primers are included in Supplementary Table 6. Only the mice with single *Fkbp* copy (Supplementary Table 7) were used for further mating.

### mESCs culture and establishment of *Gabpa*^dTAG/dTAG^ cell line

The laboratory-maintained ES-E14 cells ^45^ were cultured on 0.1% gelatin coated plates with 2i/LIF condition. Cells were grown in DMEM (Gibco, 11960069), supplemented with 15% fetal bovine serum (FBS) (Sigma-Aldrich, F6178), 2 mM GlutaMAX (Gibco, 35050061), 1 mM sodium pyruvate (Gibco, 11360), 1× MEM NEAA (Gibco, 11140050), 0.084 mM 2-mercaptoethanol (Gibco, 21985023), 1 mM sodium pyruvate (Gibco, 11360070), 100 U/ml penicillin-streptomycin (Gibco, 15140122), 1000 IU/ml LIF (Millipore, ESG1107), 0.5 μM PD0325901 (Tocris, 4192) and 3 μM CHIR99021 (Tocris, 4423).

To establish *Gabpa*^dTAG/dTAG^ cell line, *Gabpa*-HAL-FKBP^F36V^-2xHA-HAR and px330 were transfected into mESCs with Lipofectamine™ 2000 (Thermo, 11668030). 24 hrs later, cells were selected with puromycin (Gibco, A1113803) for another 48 hrs. Then cells were cultured in puromycin-free medium for one week. Single clones were picked for genotyping and further analysis.

### *In vitro* fertilization and embryo culture

Female mice (7-8 weeks) were superovulated through an initial injection of 7.5 IU pregnant mare serum gonadotropin (PMSG, BioVendor, RP1782725000), followed 48 hrs later with a 7.5 IU injection of human chorionic gonadotropin (hCG, Sigma, C1063). Oocyte-cumulus complexes (OCCs) were collected 14 hrs post hCG injection. Sperm was harvested from the cauda epididymis of adult male mice (8-12 weeks) 1 h before OCCs collection. The sperm suspension was capacitated for 1 h in 200 μl HTF medium (Millipore, MR-070-D). Subsequently, OCCs were exposed to spermatozoa for a 6-hrs incubation. The time when sperm were added to OCCs was considered as 0 hpf. Two-nuclear zygotes were cultured in the KSOM medium (Millipore, MR-106-D) under a humidified atmosphere of 5% CO_2_ at 37°C for further development.

### Western blot

1.5×10^6^ cells were lysed in 100 µl RIPA lysis buffer and incubated on ice for 30 mins. 90 µl supernatant were mixed with 12.5 µl 5× loading buffer, and heated at 98 °C for 15 min. Samples were run on NuPAGE 4-12% gel (Invitrogen, NP0322BOX) and transferred onto PVDF Transfer Membrane. Primary antibodies used included anti-GABPA (1:4000, Proteintech), anti-HA (1:1000, CST), and anti-β-Actin (1:5000, CST, #4967). Secondary antibodies used included Goat anti-Rabbit IgG (H+L) Superclonal™ Secondary Antibody-HRP (Thermo Scientific, A27036, 1:2000) and Goat anti-Mouse IgG (H+L) Secondary Antibody-HRP (Thermo Fisher Scientific, 31430, 1:2000). Protein bands were detected with ECL kit (Thermo Fisher Scientific, 32209) and imaged by Tanon 4600SF Imaging System (Tanon).

### Immunostaining and confocal microscope

Embryos were fixed with 4% paraformaldehyde/0.5% triton for 20 min, followed by three times of washing with PBS/0.1% triton, then blocked in PBS/1% BSA/0.01% triton for 1 h. Embryos were incubated overnight at 4 °C with primary antibodies: anti-GABPA (1:200, Proteintech, 21542-1-AP, Lot#00018047), anti-HA (1:200, CST, 2367S), anti-GATA4 (1:200, R&D Systems, MAB2606-SP), anti-NANOG (1:200, Abcam, ab80892) and anti-CDX2 (1:500, R&D Systems, AF3665-SP). Secondary antibodies used included Donkey anti-Goat IgG (H+L) Secondary Antibody, Alexa Fluor 647 (Thermo Scientific, A-21447, 1:500), Donkey anti Rabbit IgG (H+L) Secondary Antibody, Alexa Fluor 488 (Thermo Scientific, A-21206, 1:500) and Donkey anti Mouse IgG Secondary Antibody, Alexa Fluor 568 (Fisher Scientific, A10037, 1:500). After three times of washing, the embryos were incubated with secondary antibodies at room temperature (RT). DNA was stained with 10 μg/ml Hoechst 33342 (Sigma). The confocal microscope (Zeiss, LSM800) was used for fluorescence detecting.

### dTAG13 treatment

dTAG13 (Tocris, 6605) was reconstituted in DMSO to a 5 mM stock. For *Gabpa*^dTAG/dTAG^ mESCs treatment, dTAG13 was dilute in mESCs culture medium to 0.5 μM. For *Gabpa*^dTAG/dTAG^ embryos treatment, dTAG13 was dilute in KSOM to 1 μM. Embryos were washed with KSOM with dTAG13 for at least three times, then culture in KSOM with dTAG13 for further development.

### CUT&RUN, ATAC and RNA-seq library preparation and sequencing

For mESCs CUT&RUN with more than 10K, cells were resuspended in 50 μl washing buffer (20 mM HEPES/pH=7.5, 150 mM NaCl, 0.5 mM spermidine and 1× protease inhibitor) with activated Concanavalin A Magnetic Beads (Polysciences, 86057-3) for 10 mins at RT, then samples were incubated with anti-GABPA (1:40, Proteintech, 21542-1-AP, lot#00018047. Note, this is the only lot that worked in CUT&RUN in our hands) overnight at 4°C. For low input mESCs and embryo CUT&RUN, some modifications were made. Briefly, mESCs, zona-free embryos or isolated ICM were resuspended in 50 μl washing buffer with activated Concanavalin A Magnetic Beads for 10 mins at RT, then samples were fixed with 1% formaldehyde for 1 min. After three times of wash with washing buffer, samples were incubated with anti-GABPA overnight at 4°C. Samples were incubated with 2.8 ng/μl pA-MNase (home-made) for 2 hrs at 4 °C. Subsequently, samples were incubated with 200 μl pre-cooled 0.5 μM CaCl_2_ for 20 mins at 4 °C and quench by adding 23 μl 10×stop buffer (1700 mM NaCl, 20 mM EGTA, 100 mM EDTA, 0.02% Digitonin, 250 µg/ml glycogen and 250 µg/ml RNase A). DNA Fragments were released by incubation at 37°C for 15 mins. For both fixed and unfixed cells, 2.5 μl 10% SDS and 2.5 μl 20 mg/ml Protease K (Thermo Fisher) was added and incubated at 55 °C for at least 1 h for reverse crosslinking. DNA was extracted by phenol-chloroform followed by ethanol precipitation. Subsequent procedure is the same as described above. Sequencing libraries were prepared with NEBNext Ultra II DNA library preparation kit for Illumina (New England Biolabs, E7645S).

ATAC-seq was performed as previously described with some modifications ^46^. Briefly, ESCs and isolated ICM were digested with adapter-loaded Tn5 for 15 mins at 37°C, and stopped by stop buffer (100 mM Tris/pH=8.0, 100 mM NaCl, 40 µg/ml Proteinase K and 0.4% SDS) and incubated overnight at 55°C. 5 µl of 25% Tween-20 was added to quench SDS. Sequencing libraries were prepared with NEBNext High-Fidelity 2×PCR Master Mix (NEB, M0541S).

For RNA-seq, fresh ESCs or embryos were collected. Reverse transcription and cDNA amplification were performed with SMART-Seq™ v4 Ultra™ Low Input RNA Kit (Clontech, 634890), followed by cDNA fragmentation, adaptor ligation, and amplification using Nextera® XT DNA Sample Preparation Kit kit (Illumina, FC-131-1024).

All libraries were sequenced by NextSeq 550 system (Illumina) with paired-ended 75-bp reads (Supplementary Table 8).

### Immunosurgery

ICMs were isolated as previously described ^47^. Briefly, blastocysts at E3.5 or E4.5 stages were collected by removing the zona pellucida with Acidic Tyrode’s solution (Millipore). Embryos were then treated with anti-mouse serum antibody (Sigma-Aldrich, M5774-2ML, 1:5 dilution in KSOM) for 30 min at 37 °C. After washed for three times with KSOM, embryos were treated with guinea pig complement (Millipore, 1:5 dilution in KSOM) for another 20 min at 37 °C. Then, the trophectoderm cells were removed by a glass pipette (the inner diameter is around 40-50 μm). E4.5 ICM refers to the mixture of EPI and PrE cells after removing TE cells with immunosurgery.

### RNA-seq data analysis

The raw sequencing reads were trimmed with Trimmomatic ^48^ (v0.39) to remove sequencing adaptors. Then, the reads were mapped to GRCm38 genome using STAR ^49^ (v2.7.8a). Gene expression levels were quantified with RSEM ^50^ (v1.3.1). To identify differentially expression genes, DESeq2 ^51^ (v1.32.0) package in R was used. The significantly differential expressed genes were called with an adjusted P-value cutoff of 0.05, fold change cutoff of 2, and mean FPKM cutoff of 1. GO enrichment was performed using R package clusterProfiler ^52^. GSEA analysis was performed using R clusterProfiler ^52^ and enrichplot.

### CUT&RUN data analysis

The raw reads were trimmed with Trimmomatic ^48^ (v0.39) to remove sequencing adaptors then mapped to GRCm38 reference genome using bowtie2 ^53^ (v2.4.2). PCR duplicates were removed with Picard MarkDuplicates (v2.23.4). Reads with mapping quality less than 30 were removed. The mapped reads were further filtered to only retain proper paired reads with fragment length between 10 and 120. Peaks were called with MACS2 ^54^ (v2.2.7.1). Reproducible peaks were generated with the IDR framework ^55^ using two replicates, with IDR threshold of 0.05. For E4.5 ICM and ESCs, we further filtered the peaks to keep the ones with q-value ≤ 10^-30^ to remove weak peaks. The signal tracks were generated with deeptools ^56^ bamCoverage (v3.5.1) with bin size of 1 and normalized by CPM. For z-score normalized signal tracks, we first used bamCoverage with bin size of 100 to generate FPKM signals then used a customized script to calculate the z-score of each bin. For late 2-cell and 8-cell GABPA ultra-low-input CUT&RUN data, we noticed low mapping rates of the raw data. Further examination of the unmapped reads suggested they were environmental DNA from human, bacteria, vectors etc. and due to the ultra-low-input cells and small number of GABPA binding regions, the ratio of mapped reads from GABPA bound DNA versus unmapped reads arising from environmental DNA were low. But this would not affect the identification of GABPA peaks, since 1) the discarded reads were unmappable to mouse reference genome, and 2) upon GABPA degradation these peaks disappeared.

The peaks were annotated with R package ChIPseeker ^57^. Peaks within −1000 to +500 around TSS were considered as promoter peaks.

The heatmaps of binding profiles were calculated with deeptools ^56^ computeMatrix (v3.5.1) using bigwig signal tracks as input and bin size of 10, and visualized in R with packages profileplyr and EnrichedHeatmap ^58^.

### ATAC-seq data analysis

ATAC-seq data were analyzed with the ENCODE ^59^ ATAC-seq pipeline with default parameters (v2.1.2, https://github.com/ENCODE-DCC/atac-seq-pipeline).

### Motif enrichment analysis

Motif enrichment analysis was performed with HOMER ^60^ (v4.11) findMotifsGenome.pl with mm10 reference and parameter -size 200, using peaks file as input.

### Motif occurrence analysis

Motif occurrence analysis was performed with HOMER ^60^ (v4.11) annotatePeaks.pl with parameters: “mm10 -size -2000,2000 -hist 20 -ghist” for peaks regions, and parameters: “mm10 - size -500,500 -hist 20 -ghist” for regions around genes TSS. The motif files were download from JASPAR database ^61^ and manually converted to HOMER motif format. We used the log odds detection threshold of 6.0 for all TFs we analyzed. The motif occurrence matrix was visualized in R with package EnrichedHeatmap ^58^.

The JASPAR motif IDs for the TFs we analyzed were: GABPA - MA0062.2; OBOX - PH0121.1; DUX - MA0611.1; NFYA - MA0060.1; NR5A2 - MA0505.1; OCT4 - MA1115.1; SOX2 - MA0143.1; NANOG - MA2339.1; TFAP2C - MA0524.2; GATA3 - MA0037.4.

### Identification of ICM/TE genes

ICM/TE genes were identified using bulk RNA-seq data of E3.5 ICM and E3.5 TE from Zhang et al. ^18^. DESeq2 ^51^ (v1.32.0) package in R was used to find the differential expressed genes between ICM and TE, with an adjusted P-value cutoff of 0.05 and fold change cutoff of 2. Genes with 8-cell FPKM ≥ 1 were defined as early ICM/TE genes, while genes with 8-cell FPKM < 1 were defined as late ICM/TE genes.

### Identification of EPI/PrE genes

EPI/PrE genes were identified using single-cell RNA-seq data of E4.5 embryos from Mohammed et al. ^62^. R package Seurat ^63^ (v5.0.1) function FindMarkers was used to identify the differential expressed genes between E4.5 EPI and E4.5 PrE, with an adjusted P-value cutoff of 0.05, fold change cutoff of 4 and cell expression percentage cutoff of 0.8. Genes with expression in at least 50% E3.5 single cells were considered as early EPI/PrE genes, while the others were considered as late EPI/PrE genes.

### Statistics and reproducibility

Student’s t-tests for graph analysis were performed with Microsoft Excel (2016). Individual data points were shown as dots in the figure panels involving Student’s t-test. Data distribution was assumed to be normal but this was not formally tested. For the immunofluorescence and western blot experiments, at least three independent repetitions were performed with consistent results, and representative data were presented. p values < 0.05 were considered statistically significant. No statistical methods were used to predetermine sample sizes but our sample sizes are similar or greater to those reported in previous publications ^8^. No data were excluded from the analyses. The experiments were not randomized. Data collection and analysis were not performed blind to the conditions of the experiments.

## Data Availability

Sequencing data that support the findings of this study have been deposited in the Gene Expression Omnibus (GEO) under accession code GSE263171. Public data use in this study: RNA-seq of mouse MII oocyte to 8-cell ^17^: GSE71434. RNA-seq of mouse E3.5 ICM and TE ^18^: GSE76505. RNA-seq of mouse E4.5 TE ^9^: GSE216256. scRNA-seq of mouse E4.5 EPI and PrE ^64^: GSE159030. scRNA-seq of mouse early embryos ^19^: GSE45719. scRNA-seq of mouse E3.5 and E4.5 embryos ^62^: GSE100597. ATAC-seq of mouse early embryos ^16^: GSE66390. NR5A2 binding in mouse embryos ^10^: GSE229740. TFAP2C binding in mouse embryos ^9^: GSE216256. SOX2 binding in mouse embryos ^8^: GSE203194. Source data are provided with this study. All other data supporting the findings of this study are available from the corresponding author on reasonable request.

## Supplementary Files

**Supplementary Table 1.** Gene lists used in this study.

**Supplementary Table 2.** Differential expressed genes of late 2-cell treated with GABPA dTAG13 versus that treated with DMSO.

**Supplementary Table 3.** Differential expressed genes of morula treated with GABPA dTAG13 versus that treated with DMSO.

**Supplementary Table 4.** Differential expressed genes of E4.5 ICM treated with GABPA dTAG13 versus that treated with DMSO.

**Supplementary Table 5.** Differential expressed genes of ESC treated with GABPA dTAG13 versus that treated with DMSO.

**Supplementary Table 6.** Oligos used in this study.

**Supplementary Table 7.** ddPCR validate *Gabpa*^dTAG^ knock-in copy number

**Supplementary Table 8.** Summary of the sequenced libraries in this study.

**Source Data files**

**Source Data Fig.1.** Unprocessed scans of western blots

**Source Data Statistical Source.** Statistical source data

## Extended data

**Extended Data Fig. 1.**
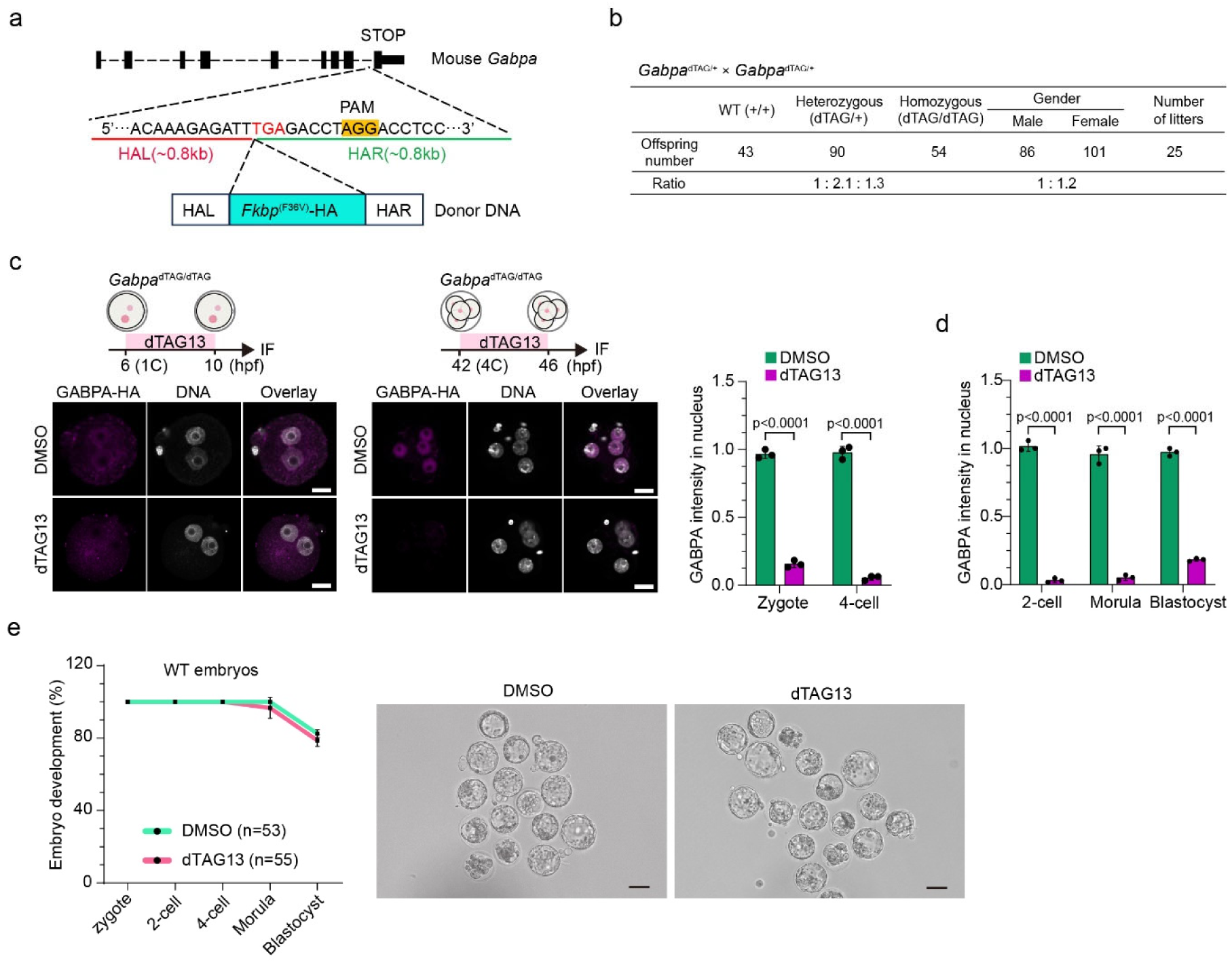
Generation of *Gabpa*^dTAG/dTAG^ mice. (**a**) Diagram of *Fkbp*^(F36V)^-HA knock-in at the 3’ end of *Gabpa* gene. (**b**) Genotypes of pups from heterozygous *Gabpa*^dTAG/+^× *Gabpa*^dTAG/+^ crosses. (**c**) Confirmation of GABPA degradation with 4h-dTAG13 treatment in zygote (left) and 4-cell (middle), and anti-HA fluorescence intensity quantification (right). Anti-HA antibody was used for GABPA-HA fusion protein staining. DNA, Hoechst 33342. Scale bar, 20 μm. (**d**) Quantification of the fluorescence intensity of GABPA in Figure 1g. (**e**) Embryo development of wild-type embryos with or without dTAG 13 treatment. The developmental rate (left) and representative pictures of E3.5 embryos (middle and right). The independent experiments were repeated three times.

**Extended Data Fig. 2.**
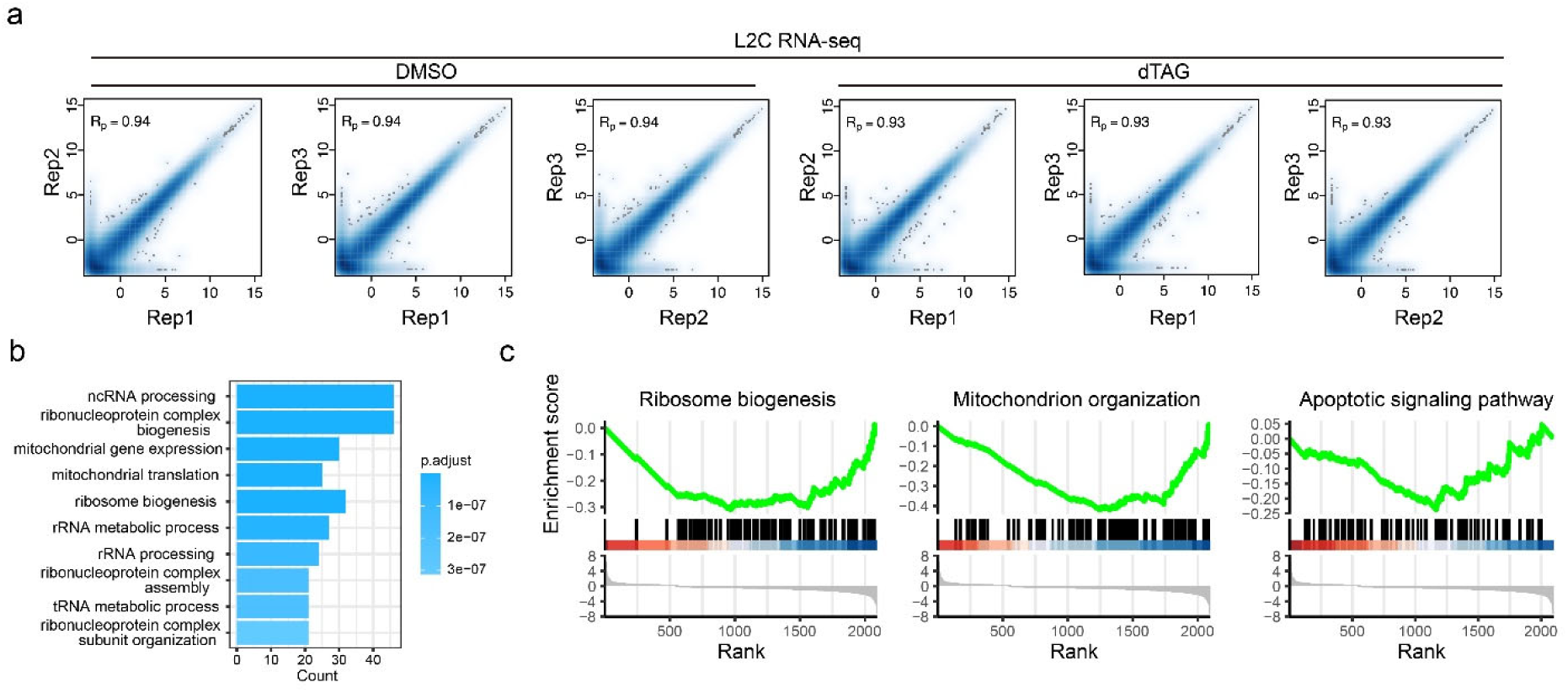
Genes affected by GABPA degradation at late 2-cell. (**a**) Correlations of the replicates of the RNA-seq at late 2-cell stage. R_p_: Pearson correlation. (**b**) Gene Ontology (GO) terms enriched for all the down-regulated genes after dTAG13 treatment at late 2-cell stage. (c) Gene set enrichment analysis (GSEA) of all the down-regulated genes at late 2-cell stage after dTAG13 treatment for ribosome biogenesis (left), mitochondrion organization (middle) and apoptotic signaling pathway (right).

**Extended Data Fig. 3.**
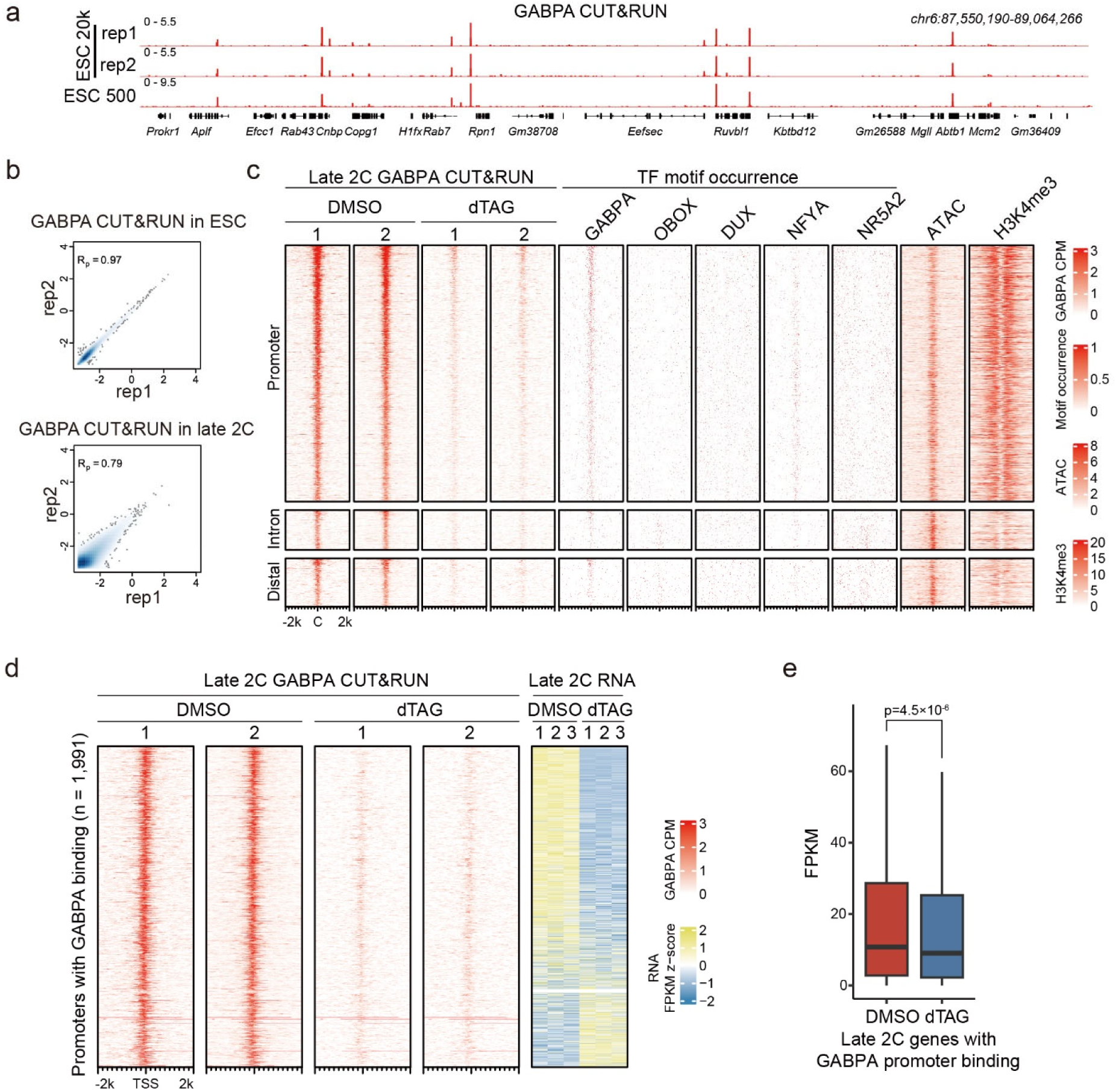
GABPA binding profiles at the late 2-cell stage. (**a**) Genome browser view showing examples of GABPA CUT&RUN profiles in ESCs using 500 and 20k cells. (**b**) Correlation of replicates for GABPA CUT&RUN with ESCs (top) and late 2-cell embryos (bottom). (**c**) Heatmaps showing the GABPA binding profiles at promoter, intron and distal regions in late 2-cell embryos. TFs motif occurrence, ATAC and H3K4me3 signals are shown at the corresponding regions. C: peak center. (**d**) Heatmaps showing GABPA promoter binding at late 2-cell stage and the corresponding genes expression level with or without the dTAG13 treatment. (**e**) Boxplot comparing the expression levels of GABPA targeted genes in DMSO and dTAG-13 treated late 2-cell. p values were calculated with two-sided Mann-Whitney U test.

**Extended Data Fig. 4.**
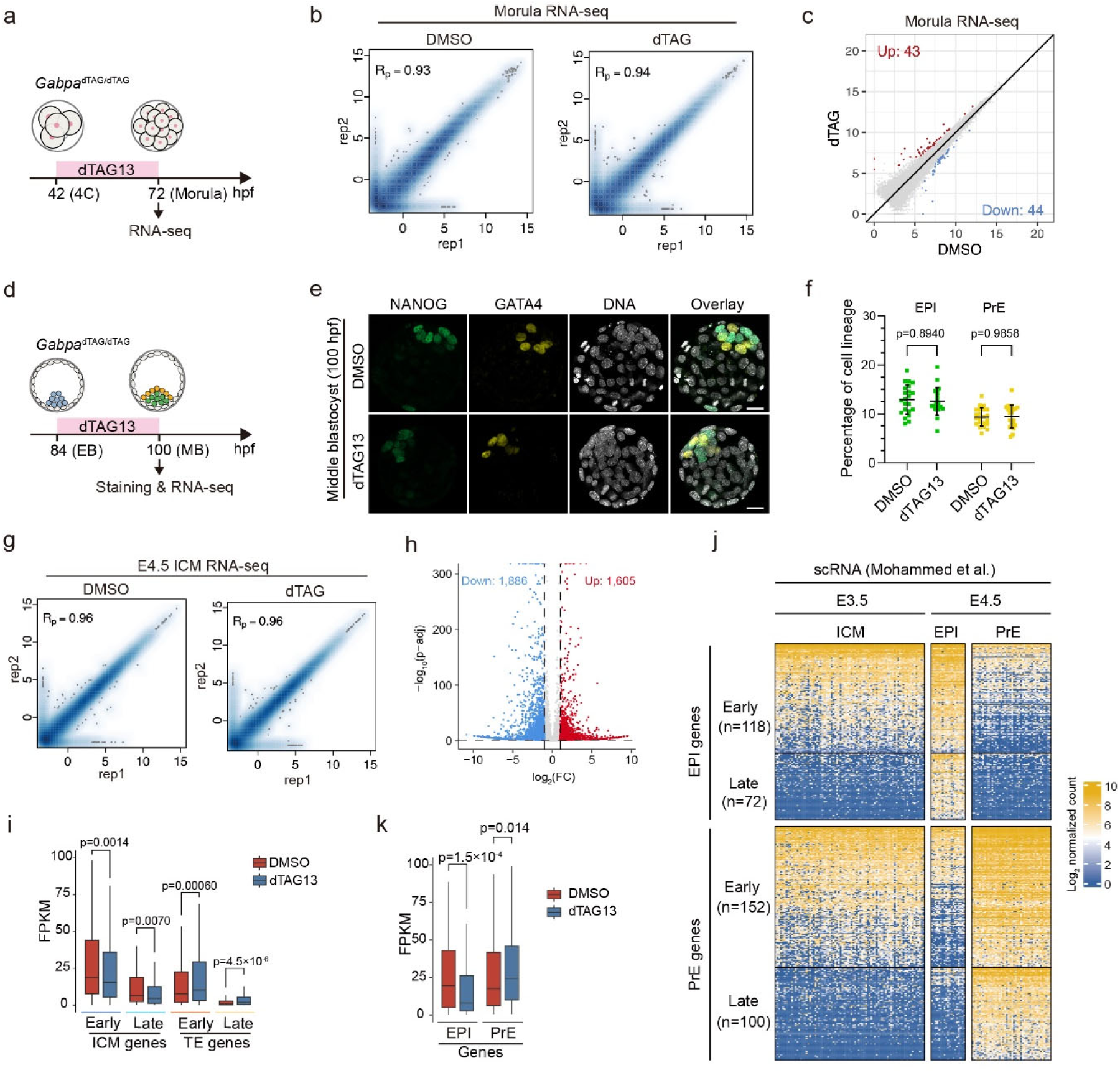
RNA-seq in morula and E4.5 ICM in response to dTAG13 treatment after major ZGA. (**a**) Diagram showing dTAG13 treatment time window and the time of sample collection for RNA-seq. (**b**) Correlation of RNA-seq replicates at morula stage. R_p_: Pearson correlation. (**c**) Dot plot showing the DEGs in morula after dTAG13 treatment shown in panel a. (d) Diagram showing the times of dTAG13 treatment and sample collection. EB: early blastocyst, MB: middle blastocyst. (**e**) Immunostaining of NANOG (green) and GATA4 (yellow) for middle blastocyst in control (DMSO) and GABPA degradation (dTAG13) conditions. (**f**) Percentage of EPI and PrE cells of middle blastocyst in control (DMSO, n=22) and GABPA degradation (dTAG13, n=21) conditions. n are the total embryo numbers used in three independent experiments. p values were calculated with Student’s t-test. (**g**) Correlation of replicates of RNA-seq in E4.5 ICM. (**h**) Volcano plot showing the DEGs after GABPA degradation in E4.5 ICM. (**i**) Boxplot comparing the expression levels of early/late ICM and early/late TE genes at E4.5 ICM with or without dTAG13 treatment. p values were calculated with two-sided Mann-Whitney U test. (**j**) EPI genes and PrE genes identified from single-cell RNA-seq data from Mohammed et al. (**k**) Boxplot comparing the expression levels of EPI and PrE genes at E4.5 ICM with or without dTAG13 treatment. p values were calculated with two-sided Mann-Whitney U test.

**Extended Data Fig. 5.**
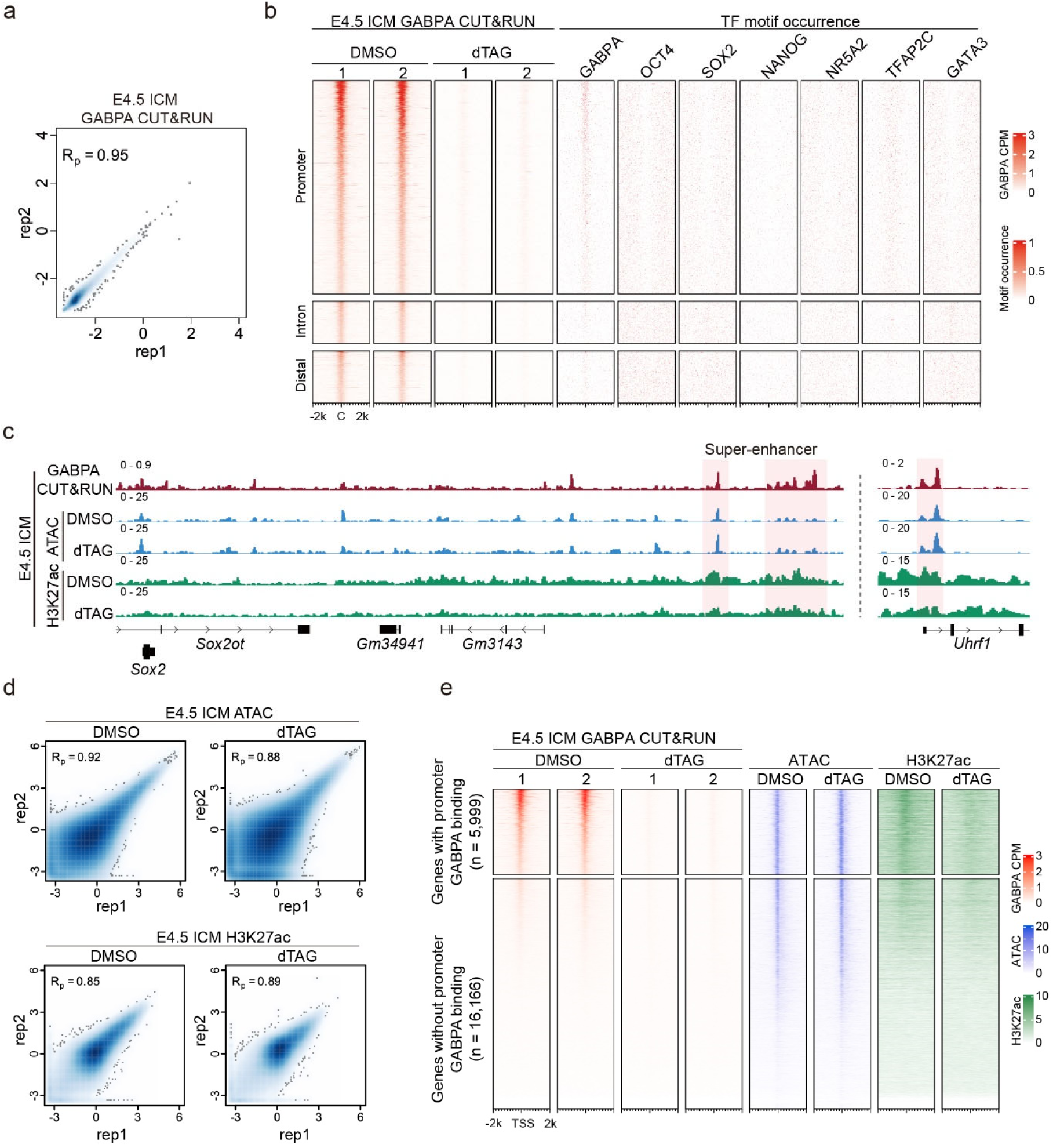
GABPA CUT&RUN in E4.5 ICM. (**a**) Correlation of replicates for GABPA CUT&RUN in E4.5 ICM. R_p_: Pearson correlation. (**b**) Heatmaps showing the GABPA binding profiles at promoter, intron and distal regions in E4.5 ICM, and the TFs motif occurrence at the corresponding regions. C: peak center. (**c**) Genome browser view of examples of GABPA binding, ATAC and H3K27ac signals at E4.5 ICM. The binding sites are shaded. (**d**) Correlation of replicates for ATAC-seq and H3K27ac CUT&RUN with or without dTAG13 treatment in E4.5 ICM. R_p_: Pearson correlation. (**e**) Heatmap showing GABPA binding, ATAC-seq and H3K27ac signals at gene promoters in E4.5 ICM. The genes were separated into two groups based on whether they have promoter GABPA binding. For ATAC and H3K27ac, two replicates in each condition were merged.

**Extended Data Fig. 6.**
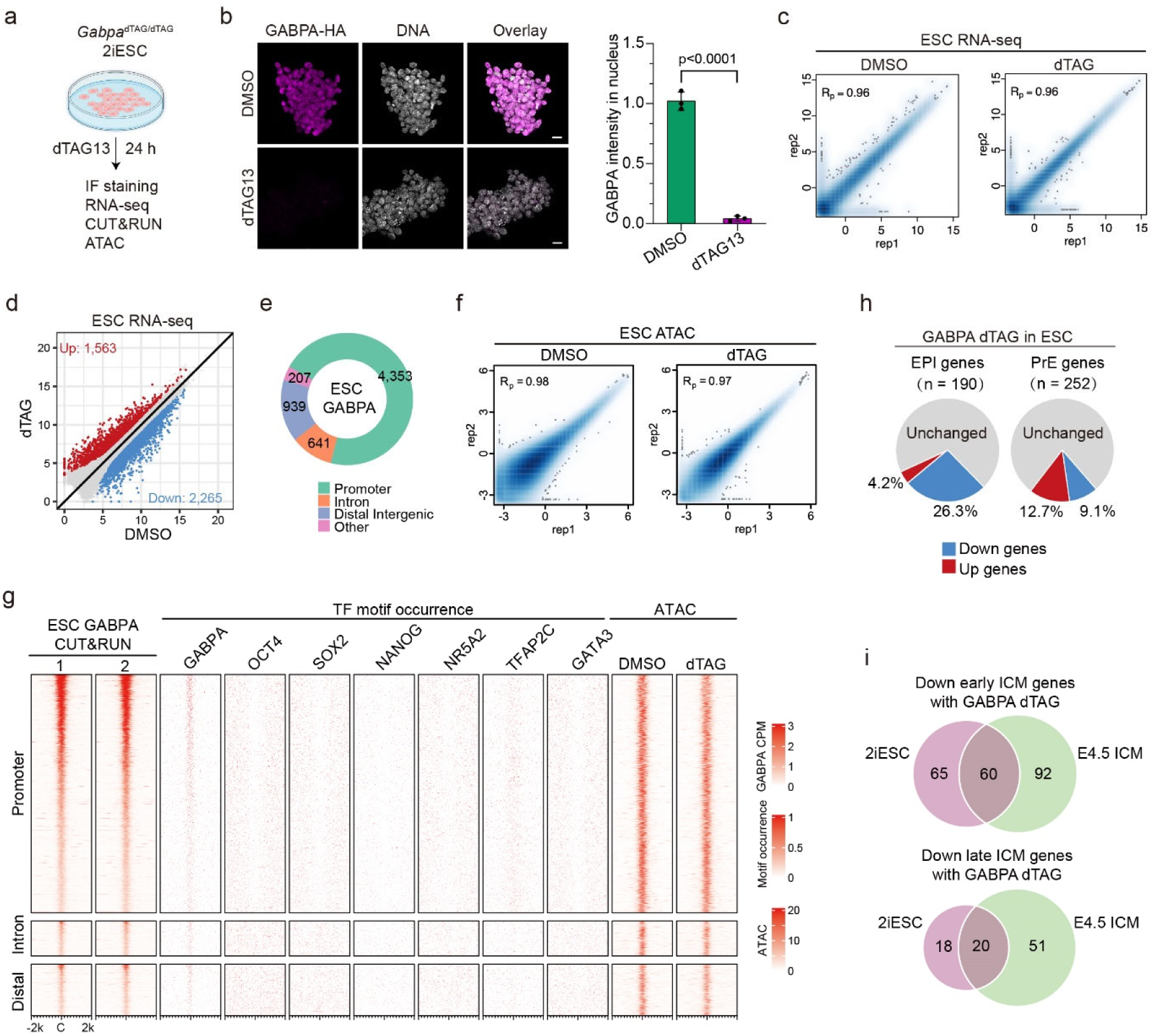
GABPA degradation in ESCs. (**a**) Diagram illustrating the GABPA degradation by dTAG13 treatment in *Gabpa*^dTAG/dTAG^ ESCs. (**b**) Immunostaining confirming GABPA degradation after dTAG13 treatment in ESCs. (**c**) Correlations of replicates RNA-seq in ESCs. R_p_: Pearson correlation. (**d**) Dot plot showing the DEGs in ESCs after GABPA degradation. (e) Genomic distribution of GABPA binding peaks in ESCs. (**f**) Correlation of replicates for ATAC-seq with or without dTAG13 treatment in ESCs. R_p_: Pearson correlation. (**g**) Heatmaps showing the GABPA binding profiles at promoter, intron and distal regions in ESCs, and the TFs motif occurrence and ATAC signals at the corresponding regions. C: peak center. (**h**) Percentage of down-regulated EPI and PrE genes after dTAG13 treatment in ESCs. (**i**) Comparisons of the down-regulated early and late ICM genes upon dTAG13 treatment in ESCs and E4.5 ICM.

**Extended Data Fig. 7.**
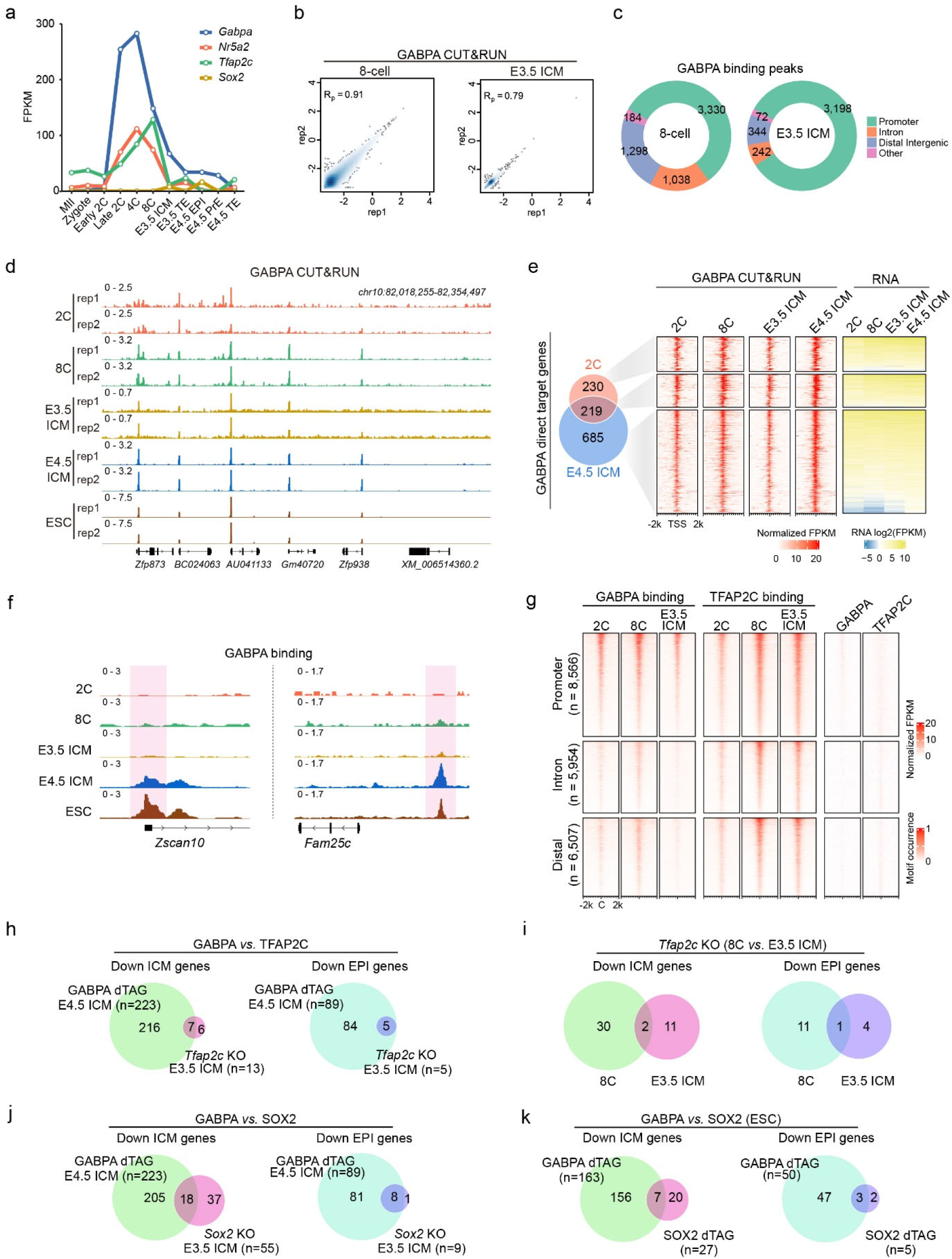
GABPA binding dynamics and comparisons with other pluripotency regulators. (**a**) Expression dynamics of *Gabpa*, *Nr5a2*, *Tfap2c* and *Sox2* during mouse preimplantation development. (**b**) Correlation of replicates for GABPA CUT&RUN in 8-cell embryos (left) and E3.5 ICM (right). R_p_: Pearson correlation. (**c**) Genomic distribution of GABPA binding peaks in 8-cell embryos (left) and E3.5 ICM (right). (**d**) Genome browser view showing examples of the binding profiles of GABPA in 2-cell, 8-cell, E3.5 ICM, E4.5 ICM and ESCs. (**e**) Comparison of the GABPA direct target genes between 2-cell embryos and E4.5 ICM. (**f**) Examples of genome browser view illustrating the GABPA binding (shaded) dynamics during preimplantation development. (g) Heatmap comparison of GABPA and TFAP2C binding profiles at promoter, intron and distal regions in 2-cell, 8-cell and E3.5 ICM. (**h**) Comparisons of GABPA (E4.5 ICM) and TFAP2C (E3.5 ICM) affected ICM genes (left) and EPI genes (right). (**i**) Comparisons of TFAP2C affected ICM genes (left) and EPI genes (right) in 8-cell and E3.5 ICM. (**j**) Comparisons of GABPA (E4.5 ICM) and SOX2 (E3.5 ICM) affected ICM genes (left) and EPI genes (right). (**k**) Comparisons of GABPA and SOX2 affected ICM genes (left) and EPI genes (right) in ESCs.

